# Two Novel Genes, *Stb23* and *Stb24*, Conferring Multi-stage Resistance to *Zymoseptoria tritici*: Rapid Deployment in Marker-Assisted Wheat Breeding

**DOI:** 10.64898/2026.04.28.717151

**Authors:** Nannan Yang, Ben Ovenden, Brad Baxter, Simon Williams, Peter S. Solomon, Andrew Milgate

**Affiliations:** Department of Primary Industries and Regional Development, Wagga Wagga Agricultural Institute, Pine Gully Road, Wagga Wagga, NSW 2650, Australia; Division of Plant Sciences, Research School of Biology, The Australian National University, Canberra 2601, Australia

## Abstract

The fungal pathogen *Zymoseptoria tritici* poses a major global threat to wheat production, causing severe yield losses and necessitating intensive and costly fungicide applications. The increasing demand for durable genetic resistance has intensified interest in quantitative resistance loci, particularly those exhibiting multi-stage resistance (MSR), which suppress pathogen development continuously throughout the wheat life cycle. Many previously effective resistance genes are now showing declining efficacy, underscoring the urgent need for novel and long-lasting sources of resistance. In this study, we report the identification and genetic mapping of two quantitative resistance loci that address this need. The first locus, designated *Stb23*, is a major QTL on chromosome 1DS, with LOD scores exceeding 9 and explaining 6-36% of phenotypic variation at the seedling stage and 2-16% at the adult-plant stage. The second locus, designated *Stb24*, is a major QTL on chromosome 3DL, with LOD scores of approximately 10 and accounting for 11-30% of seedling-stage variation and 9-23% of adult-plant variation. Furthermore, two tightly linked KASP markers-snp_1D1217527 for *Stb23* and snp_3D1077880 for *Stb24*-were developed and validated across three popular Australian bread wheat cultivars, providing practical tools for deploying these loci in breeding programs targeting improved resistance to *Z. tritici*.

**Key message:** Two significant major-effect resistance loci on chromosomes 1DS (proposed as *Stb23*) and 3DL (proposed as *Stb24*) were identified and characterized. Two tightly linked KASP markers with these loci were also discovered and validated for molecular-assisted breeding programs.

## Introduction

*Zymoseptoria tritici* (syn. *Mycosphaerella graminicola* (Fuckel) J. Schrot, anamorph *Septoria tritici*) (Quaedvlieg et al. 2011) remains one of the most damaging pathogens affecting global wheat production, particularly in Australia, Europe, and North America. It is ranked as the third most damaging pathogen affecting wheat yields worldwide, causing an estimated average annual loss of 2.44% (Savary et al. 2019). Under favourable environmental conditions-characterised by mild temperatures and extended periods of leaf wetness-yield losses can increase dramatically, often reaching 30-50% (Eyal 1987).

Of the 23 reported *Stb* resistance genes (Brown et al. 2015; Langlands-Perry et al. 2022; Yang et al. 2018), only three have been cloned to date. *Stb6*, a wall-associated receptor kinase-like gene (*TaWAKL4*), mediates a defense response that restricts pathogen penetration without triggering a hypersensitive response (Saintenac et al. 2018). *Stb16q*, encoding a cysteine-rich receptor-like kinase (CRK), acts very early in the infection cycle by halting fungal development at the stomatal entry point (Saintenac et al. 2021). The recently cloned *Stb15*, a G-type lectin receptor-like kinase (LecRLK), further expands the range of receptor-like kinase-mediated mechanisms known to underlie resistance to *Z. tritici* (Hafeez et al. 2025). Nonetheless, rapid pathogen evolution has compromised the longevity of several deployed resistance genes, including *Stb1*, *Stb4*, and *Stb16q* (Kildea et al. 2020; Ponomarenko et al. 2011). This reinforces the importance of quantitative or partial resistance, which is generally more durable against *Z. tritici* (Brown 2015; Niks et al. 2015; Rimbaud et al. 2021; Thauvin et al. 2024). Notably, no cloned *Stb* gene to date resembles the classic adult-plant “slow-rusting” genes such as *Lr34*, *Lr46*, or *Lr67*, which provide durable, partial resistance by limiting disease progression rather than preventing infection outright (Krattinger et al. 2009; Moore et al. 2015).

Consequently, the search for novel resistance sources and alternative resistance-gene architectures remains a priority. Quantitative effect loci conferring **multi-stage resistance (MSR)** are of particular interest because they reduce pathogen inoculum across developmental stages and effectively suppress epidemic development (Yang et al. 2022). According to the review by Brown et al. (2015), *Stb1*, *Stb4*, *Stb5*, and *Stb18* can be considered as MSR loci. For instance, *Stb18*, derived from the cultivar ‘Balance’, reduces disease severity by up to 48% at the seedling stage and by approximately 13% at the adult-plant stage (Tabib Ghaffary et al. 2011). Importantly, resistance expressed at the seedling stage does not always translate to adult-plant resistance, highlighting the need to evaluate resistance across multiple developmental stages in breeding programs. For example, only one of twenty QTLs (Thauvin et al. 2024) and four of thirteen QTLs captured at the seedling stage were also detected at the adult-plant stage (Yang et al. 2022).

In Australia, ten previously reported *Stb* genes-including *Stb2* from ‘Veranopolis’, *StbWW* from ‘WW2449’, *Stb3* from ‘Israel 493’, *Stb4* from ‘Tadorna’, *Stb6* from ‘Hereward’, *Stb8* from ‘W7984’, *Stb9* from ‘Soissons’, *Stb14* from ‘M1696’, *Stb15* from ‘Arina’, and *Stb18* from ‘Balance’-have shown declining effectiveness against local *Z. tritici* populations (unpublished data). This underscores the urgent need to identify and characterize resistance sources that are robust across the genetic diversity present within Australian pathogen populations. To enhance the efficiency and scalability of marker-assisted selection (MAS), reliable molecular markers are essential. Kompetitive Allele Specific PCR (KASP) markers are particularly valuable for rapid integration of resistance genes into breeding pipelines. To date, only six validated KASP markers linked to four major *Stb* genes are available: two for *Stb6* (Thauvin et al. 2024), one for *Stb15* (Hafeez et al. 2025), two for *Stb16q* (Saintenac et al. 2018), and one for *Stb19* (Yang et al. 2018).

In this study, we applied a semi-double-blind phenotyping and genotyping framework to enhance the efficiency of identifying and deploying resistance traits within advanced wheat breeding materials. Using this approach, BC₁F₂ populations, derived from resistant F₂ individuals, were screened and genotyped for field-based resistance, while their corresponding F₄ progeny were assessed and genotyped concurrently for seedling-stage resistance. Importantly, all evaluations were performed without prior knowledge of the underlying resistance loci, ensuring unbiased detection of both isolate-specific and multiple-growth-stage resistance. By coordinating phenotyping across successive developmental stages, this framework reduced the likelihood that seedling-stage resistance would fail to translate into adult-plant performance and substantially shortened the breeding cycle required to integrate newly discovered resistance loci into advanced wheat lines.

Using this approach, we successfully identified and genetically characterised two **MSR** loci effective against *Z. tritici*. The first, a major QTL on chromosome **1DS**, is designated ***Stb23*** in this study. The second, located on chromosome **3DL**, is designated ***Stb24***. Both loci conferred resistance consistently across seedling assays and multiple adult-plant growth stages, underscoring their robustness and importance for durable resistance breeding. To facilitate breeding deployment, we developed tightly linked KASP markers for both loci. These diagnostic markers enable efficient marker-assisted selection and support the incorporation of *Stb23* and *Stb24* into elite wheat germplasm, thereby enhancing resistance to *Z. tritici* across diverse environments.

## Materials and methods

### Plant Materials

Our study used 11 segregating populations, detailed in Table 1. A key aspect of this investigation involved analysing a single population across three generations. The two primary populations were X6135 (‘WW2449’ x ‘WW33612’) and X6129 (‘WW2449’ x ‘WW33611’).

**Table 1.** Segregating populations used in this study.

| Cross Name | Pedigree | Pop | Pop Size | Score Method |
| --- | --- | --- | --- | --- |
| X6135 | WW2449/WW33612 | F<sub>2</sub> | 65 | Single Plant |
| X6135 | WW2449/WW33612 | F<sub>2:4</sub> | 188 | Single Plant |
| X6135 | WW2449/WW33612 | F<sub>2:6</sub> | 153 | Replicated Rows |
| X6213 | WW2449*2/WW33612 | F<sub>2</sub> | 88 | Single Plant |
| X6284-6 | Lancer*2/WW33612 | F<sub>2</sub> | 15 | Single Plant |
| X6312-2 | Suntop*2/WW33612 | F<sub>2</sub> | 25 | Single Plant |
| X6312-10 | Suntop*2/WW33612 | F<sub>2</sub> | 29 | Single Plant |
| X6129 | WW2449/WW33611 | F<sub>2</sub> | 95 | Single Plant |
| X6129 | WW2449/WW33611 | F<sub>2:4</sub> | 188 | Single Plant |
| X6129 | WW2449/WW33611 | F<sub>2:6</sub> | 149 | Replicated Rows |
| X6211 | WW2449*2/WW33611 | F<sub>2</sub> | 90 | Single Plant |
| X6285-3 | Lancer*2/WW33611 | F<sub>2</sub> | 15 | Single Plant |
| X6285-5 | Lancer*2/WW33611 | F<sub>2</sub> | 38 | Single Plant |
| X6293-5 | Scepter*2/WW33611 | F<sub>2</sub> | 27 | Single Plant |
| X6300-1 | Suntop*2/WW33611 | F<sub>2</sub> | 35 | Single Plant |

‘WW2449’ is a spring wheat cultivar exhibiting high susceptibility to *Z. tritici* at both seedling and adult-plant stages. Despite this, it possesses the *StbWW* gene (Table 2), which confers resistance to the WAI332 isolate when inoculum was sprayed to seedlings (Raman et al., 2009). ‘WW33612’ is a Single Seed Descent (SSD) line, selected for resistance to *Z. tritici* in Australia, originating from the resistant synthetic cultivar ‘W7984’. This line consistently demonstrates stable resistance to *Z. tritici* at both seedling and adult-plant stages. Prior research indicates that ‘W7984’ carries the adult-plant resistance gene *Stb8*, effective against the IN95-Lafayette-1196-WW 1-4 isolate (Adhikari et al. 2003). Similarly, ‘WW33611’ is an SSD line selected for resistance to *Z. tritici* in Australia, derived from the resistant cultivar ‘Fleche D’or’ originating from France (Ballantyne 1989), also showing stable resistance at both seedling and adult-plant stages.

**Table 2.**
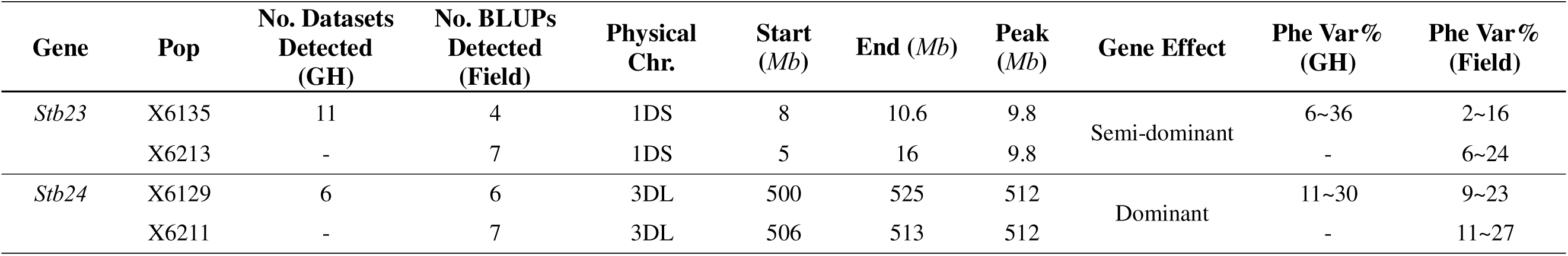
Summary of the *Stb23* and *Stb24* loci on the IWGSC CS RefSeq v2.1 physical map, including their physical positions, peak intervals, gene effects, and phenotypic contributions (Phe Var%) detected in greenhouse and field trials across four wheat segregating populations.

To validate our findings, nine BC_1_F_2_ segregating populations were generated (Table 1). These populations were created using ‘WW2449’, alongside STB-susceptible cultivars ‘Lancer’, ‘Scepter’, and ‘Suntop’ which are widely utilized as parents in Australian breeding programs.

#### Glasshouse Screening for *Zymoseptoria tritici* Resistance

We utilized three Australian *Z. tritici* isolates for our glasshouse experiments: WAI332 (collected from NSW in 1979), WAI251 (from Victoria in 2012), and WAI161 (from Tasmania in 2011). The inoculation procedure, involving an infiltration method for phenotyping *Z. tritici* glasshouse infections, detailed in previously described methods (Yang et al. 2018). All three isolates (WAI332, WAI251, WAI161) were tested on X6135 F_2_ and X6129 F_2_ populations (Supplementary Table 1 and 2). Additionally, isolates WAI161 and WAI332 were tested on X6135 F_4_, and WAI161 was tested on X6129 F_4_, in four independent glasshouse experiments (Figure 1).

**Fig. 1.**
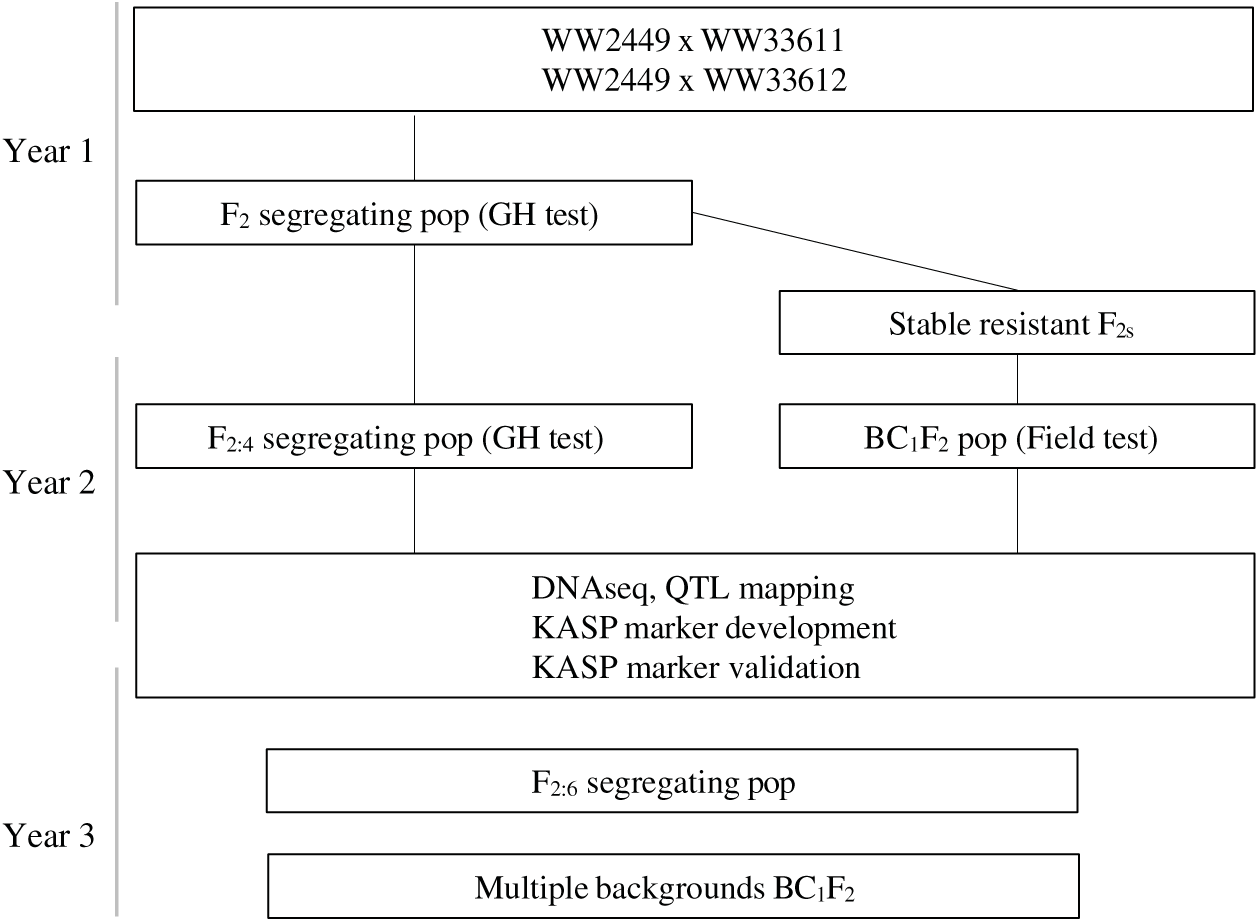
Schematic workflow of segregating populations’ development, QTL mapping, gene discovery, KASP marker development and validation, and application in wheat breeding.

*Z. tritici* symptoms were evaluated between 20 and 35 days post-inoculation. A seedling infection score (STB_S, 1-5 scale) was assigned based on a visual estimation of the percentage of leaf area exhibiting necrosis (Nec%, 0-100%). Additionally, pycnidia density on the necrotic leaf area (Pyc%, 0-100%) was recorded at multiple time points after inoculation. Further details of the scoring procedure are described in Yang et al. (2022).

#### Field Experiments

Field evaluations were conducted at two sites: Wagga Wagga Agricultural Institute, NSW, Australia (WGA, -35.04419222, 147.3167896) in 2023 and 2024, and Agriculture Victoria Hamilton Centre, Hamilton, Victoria, Australia (HLT, -37.828768, 142.082319) in 2024. In 2023, the X6213 F_2_ and X6211 F_2_ populations were assessed at Wagga Wagga. For 2024, we evaluated nine segregating populations: X6135 F_6_, X6129 F_6_, X6284-6 F_2_, X6312-2 F_2_, X6312-10 F_2_, X6285-3 F_2_, X6285-5 F_2_, X6293-5 F_2_, and X6300-1 F_2_. At Wagga Wagga, natural *Z. tritici* infection was augmented with stubble debris from a local crop with high levels of *Z. tritici* infection. While field experiments at Hamilton relied solely on natural infections.

Relative maturity was assessed using the Zadoks decimal growth scale (Zadoks et al. 1974) concurrently with field disease evaluations. Disease severity was visually scored using the Saari and Prescott scale (Saari and Prescott 1975). The adult-plant severity score, STB_A (1-9), quantified *Z. tritici* infection severity while accounting for the plant growth stage. The percentage of diseased leaf area (STB%, 1-100%) represented the proportion of total leaf area exhibiting necrosis or chlorosis, whereas pycnidia density (Pyc%, 0-100%) was scored on the necrotic areas of recently infected leaves. Under field conditions, the infection cycle of *Z. tritici* typically requires 28-40 days before symptoms become visible and scorable. For example, when plants reach Zadoks growth stage 51 (flag leaf fully emerged), pycnidia assessments are made on the Flag−1 and/or Flag−2 leaves. At this stage, these leaves have generally been exposed to inoculum for one or two infection cycles. Infection progression may be slower in June and July under Australian winter conditions or during extended dry periods, which can delay symptom development in the field.

For X6213 and X6211 at Wagga Wagga in 2023, three phenotypic scores were collected approximately four weeks apart in August, September, and October. In 2024, two phenotypic scores were collected for X6135 F_6_ and X6129 F_6_ at Wagga Wagga, similarly four weeks apart in September and October. Additionally, two phenotypic scores were collected for X6284-6 F_2_, X6312-2 F_2_, X6312-10 F_2_, X6285-3 F_2_, X6285-5 F_2_, X6293-5 F_2_, and X6300-1 F_2_ in September and October (Table 1).

#### Experimental Design

Spatially optimised randomised complete block designs were generated for all experiments with F_6_ progenies, using the software package DiGGer (Coombes 2002) in R (R Core Team 2020). In the 2024 field experiments, X6135 F_6_ and X6129 F_6_ progenies were replicated three times, with the susceptible control cultivar ‘WW425’ comprising the remaining entries. Spatial designs for field experiments were configured as Row x Column arrays at both Wagga Wagga, NSW, and Hamilton, VIC. However, only single plant/genotype for the F_2_ and F_4_ populations in Table 1 was assessed for its phenotypic response at multiple times in the glasshouse or field conditions, generating various datasets used for QTL analysis.

#### Phenotypic Data Analysis

For the X6135 F_6_ and X6129 F_6_ populations, data for each trait within each experiment were modelled using the software package ASReml-R 4.2 (VSNi) in R version 4.5.0 (R Core Team, 2020) following the linear mixed model approach of Gilmour et al. (1997). The models used are detailed in Yang et al. (2022). Best Linear Unbiased Predictions (BLUPs) from the models were used for subsequent linkage analyses. The model contains the experimental blocking structures (replicate, range and row) used to generate six BLUPs for the X6135 F_6_ population in the field, and nine BLUPs for the X6129 F_6_ population in the field. Details about the model used in ASReml were described in the previous study (Yang et al. 2022).

Correlation analysis among various traits was performed using R/corrplot (Wei et al. 2017), with significant differences between each pair of traits calculated using the Spearman correlation method. To compare differences among various groups of alleles, a multiple comparison Wilcoxon test was used to generate *p-value*s using R/rstatix (Kassambara 2025).

#### DNA Extraction and Genotyping

Leaf tissue from 14-day-old seedlings of X6135 F_4_ and X6129 F_4_ progenies was harvested, freeze-dried, and used for DNA extraction. Cost-reduction measures were implemented by submitting only 50% of the F_4_ progenies from the X6135 and X6129 populations for genomic analysis. Genotyping-by-Sequencing (GBS) was performed by DArT Pty Ltd (Canberra, Australia. Marker genetic positions were assigned based on IWGSC CS RefSeq v2.1, as supplied by DArT. DNA from the AuSTB panel, comprising diverse local and international cultivars, was also used for KASP marker development; the panel’s composition is detailed in our previous study (Yang et al. 2022). For X6284-6 F_2_, X6312-2 F_2_, X6312-10 F_2_, X6285-3 F_2_, X6285-5 F_2_, X6293-5 F_2_, and X6300-1 F_2_, crude DNA was extracted in early-August 2024 using Platinum™ Direct PCR Universal Master Mix (Thermo Fisher Scientific, cat#A44647500). This crude DNA was then used for KASP genotyping to select homozygous lines from these populations.

DNA samples from X6135 F_4_, X6129 F_4_, X6213 F_2_, and X6211 F_2_ were sequenced, and SNPs were called against IWGSC CS RefSeq v2.1 by DArT Pty Ltd, Canberra, Australia. Quality control revealed a reproducibility rate of 0.9 for SNPs and a call rate of 0.85 for SNPs. The experimental procedure for SNP generation is comprehensively described by (Courtois et al. 2013) and (Li et al. 2015). The resulting SNP datasets were subsequently used for linkage analysis and QTL mapping.

#### Primer Design and KASP Assays

To design primers for Kompetitive Allele-Specific PCR (KASP) assays, nine SNPs from X6135 and twelve SNPs from X6129 F_4_ were chosen for KASP development. KASP primer design was conducted using PolyMarker (Ramirez-Gonzalez et al. 2015), and the generated primers were used for genotyping of the whole population of X6135 F_4_/F_6_, X6129 F_4_/F_6_, the AuSTB panel, X6284-6 F_2_, X6312-2 F_2_, X6312-10 F_2_, X6285-3 F_2_, X6285-5 F_2_, X6293-5 F_2_, and X6300-1 F_2_.

KASP assays were performed in 96-well PCR plates, with a final reaction volume of 10 μl containing 5 μl of KASP 2x reaction mix (LGC Genomics, UK) and 0.14 μl assay mix. Further details are described in Yang et al. (2018). PCR products were detected using a FLUOstar Omega plate reader (BMG LABTECH GmbH, Ortenberg, Germany), and the resulting data were analyzed manually with Klustercaller software (version 3.4.1; LGC Hoddesdon, UK).

#### Linkage Map Construction

Given the high-density SNP coverage, the physical maps aligned to the IWGSC CS RefSeq v2.1 genome for the X6135 F₄, X6213 F₂, and X6211 F₂ populations were directly suitable for QTL analysis. The geno.table() function in the R/qtl package (Broman and Sen 2009) was used to identify and remove highly distorted SNP markers that could interfere with downstream analyses. Linkage map construction was required only for the X6129 F₄ population because its SNP order did not correspond appropriately to the reference genome for direct QTL mapping. Genotypic data were prepared using R/qtl (Broman and Sen 2009) and the ASMap package (Taylor et al. 2015), with linkage groups formed using a LOD threshold of 12 and a maximum recombination fraction of 0.35. Linkage map construction was therefore performed exclusively for the X6129 F₄ population to ensure accurate marker ordering, while the remaining populations relied directly on the IWGSC CS RefSeq v2.1 physical maps for subsequent QTL analyses.

#### QTL Mapping

The R package R/qtl (Broman and Sen 2009) was used for QTL mapping. Specifically, QTL analysis using the scanone() function (with either “mr” or “hk” method) identified potential QTLs across all datasets/BLUPs generated from the X6135, X6129, X6213, and X6211 populations. A LOD score of 2.5 was used to call a QTL. The representative marker for each QTL exceeding this threshold was then utilized for multiple interval mapping, employing functions such as ‘makeqtl’, ‘fitqtl’, and ‘refineqtl’ for all the X6135, X6129, X6213, and X6211 segregating populations (Table 1). Results reported phenotypic variances, additive effects, and evaluated potential dominance effects.

#### Bioinformatic Analysis

The QTLs identified in this study were compared to 23 published *Stb* genes and over 100 previously reported QTLs summarised in (Yang et al. 2022). First, genomic DNA sequences corresponding to the identified QTL regions were extracted from IWGSC CS RefSeq v2.1 genome. Published QTLs that spatially overlapped with our identified QTL regions were considered to co-localize. Subsequently, all Coding Sequences (CDS) were extracted from the QTL regions and subjected to BLAST analysis against the NCBI database using OmicsBox 3.4 (BioBam Bioinformatics) to annotate candidate R genes within the QTLs reported in this study.

## Results

### Phenotypic data of Segregating Populations

The experimental workflow-including population development, phenotyping strategies, and genetic analyses-is summarised schematically in Fig. 1. To maximise efficiency, we implemented a semi-double-blind phenotyping and genotyping design. In brief, BC₁F₂ populations, each derived from resistant individuals identified in the F₂ generation (Supplementary Table S1 and S2), were evaluated for field-based resistance, while their corresponding F₄ descendants were assessed for seedling-stage resistance, and the F₆ descendants were evaluated for adult-plant resistance as a cross-validation step for the BC₁F₂ adult-plant responses. This multistage assessment was conducted without prior knowledge of the underlying resistance loci, enabling unbiased detection of both isolate-specific and multiple-growth-stage resistance loci.

The evaluated traits encompassed necrotic leaf area percentage (Nec%), pycnidia density (Pyc%), and the seedling-stage STB severity scale (STB_S), as well as adult-plant traits including the STB severity scale (STB_A) and percentage of infected leaf area (STB%) (Fig. 2, Supplementary Tables S1-S2). For the X6135 population, strong positive correlations were observed between STB_S and STB_A, ranging from 0.16 to 0.93 (Fig. 3a). Likewise, Pyc% exhibited strong cross-stage correlations between seedling and adult-plant assessments, varying from 0.12 to 0.96 (Fig. 3a). In contrast, for the X6129 population the data displayed weak positive correlations between STB_S and STB_A, with values just above zero, while Pyc% correlations across growth stages ranged from 0.10 to 0.42 (Fig. 3b). Overall, the phenotypic analyses highlight the predictive value of seedling-stage responses for subsequent adult-plant resistance and demonstrate the interconnected behaviour of STB-related traits across multiple developmental stages.

**Fig. 2.**
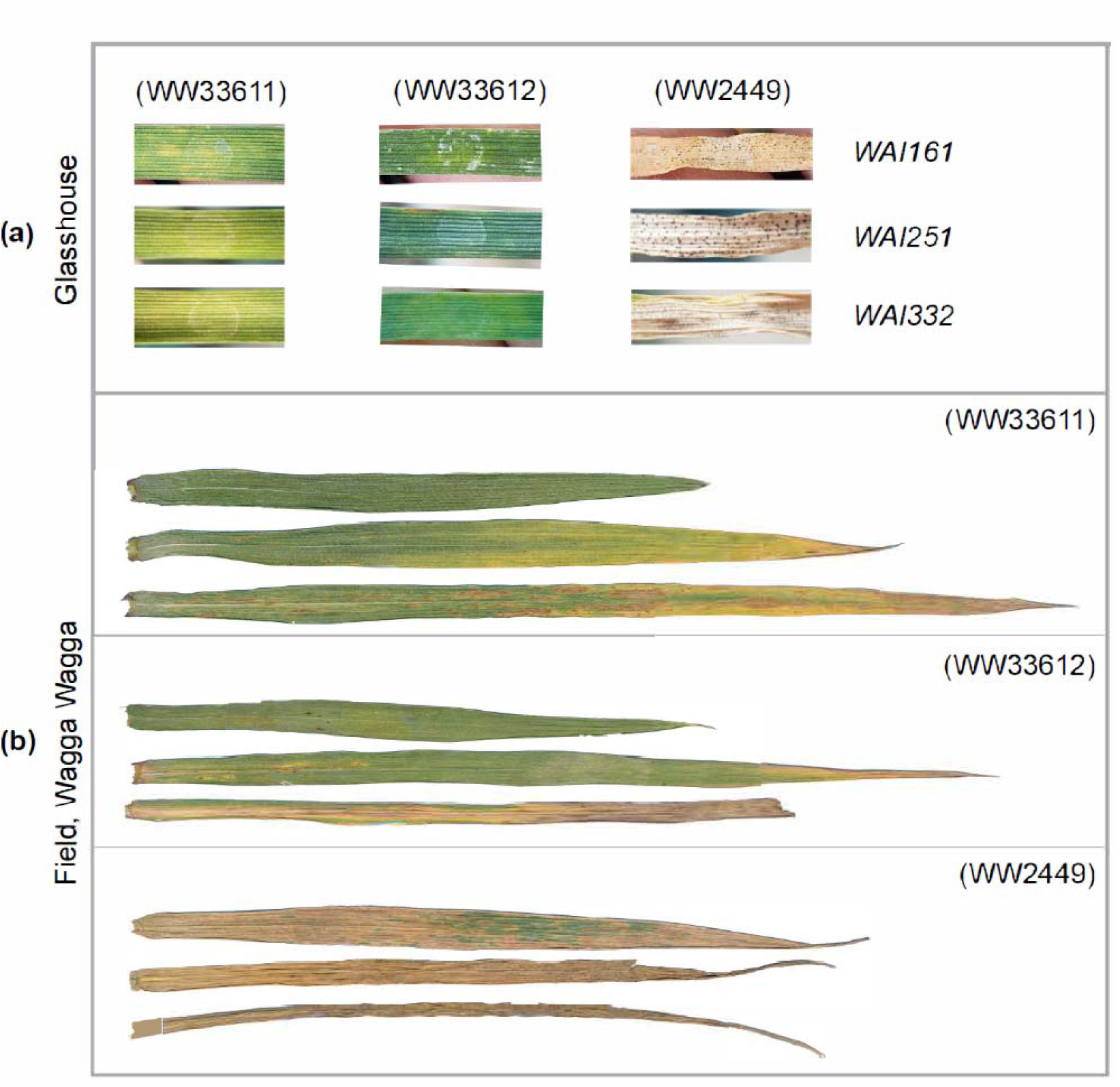
Comparisons of infected leaves between ‘WW33611’, ‘WW33612’, and ‘WW2449’ under different treatments. **(a)** Comparisons of infected leaves between ‘WW33611’, ‘WW33612’, and ‘WW2449’, infiltrated by WAI161, WAI251, and WAI332, 25 days after infection at 2-leaf seedling stage. **(b)** Comparisons of infected Flag, Flag-1, and Flag-2 leaves between ‘WW33611’, ‘WW33612’, and ‘WW2449’ at adult-plant stage Wagga Wagga NSW 2024 (WGA24).

**Fig. 3.**
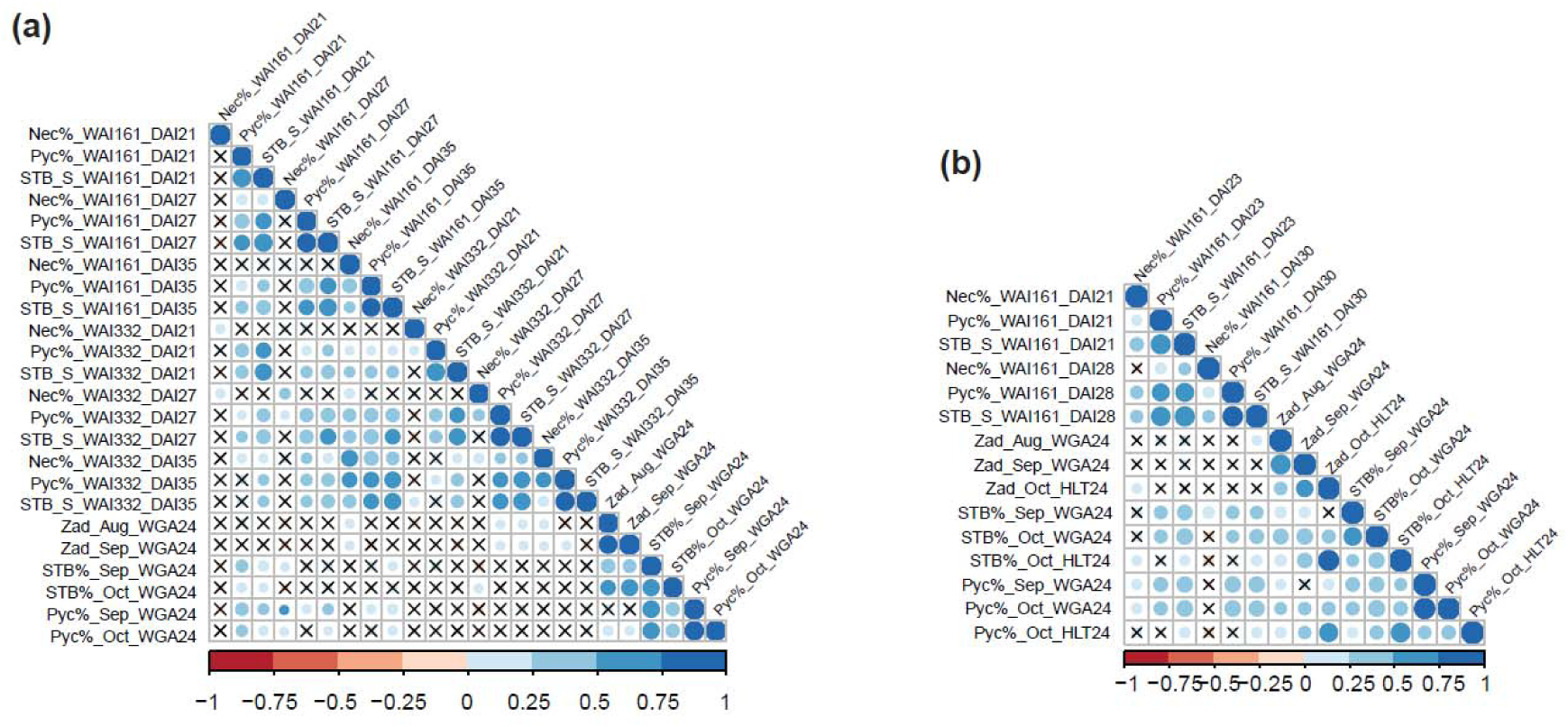
Correlation analysis between various traits measured on populations X6135 **(a)** and X6129 **(b)**. Correlation analysis between three traits measured at 2-leaf seedling stage and three traits measured at adult-plant stage. Traits at the seedling stage include the percentage of necrotic leaf area (Nec%) on the infected leaves, the pycnidia density (Pyc%) in the necrotic leaf area, and the STB_S Scale 1 to 5. Traits were measured at least 21 days after inoculation, using two STB isolates WAI161 and WAI332. Traits at the adult-plant stage include relative maturity (Zadoks scale), STB_A scale 1-9, and the percentage of STB infected leaf area on the whole plant (STB%). Traits from adult-plant stage were measured at Wagga Wagga NSW (WGA) and Hamilton Vic (HLT) in 2024. Red solid circle represents the negative correlation between traits, while blue solid circle represents the positive correlation between traits. Cross mark represents that the correlation between the traits is not significant. The darker the color, the stronger the correlation.

### Genotypic Data and Marker Distribution

Genotypic analysis was conducted to characterise the distribution and density of polymorphic SNP markers across the chromosomes of the four mapping populations, thereby providing insight into their genetic diversity and genome-wide marker coverage. The full marker distributions for the X6135, X6129, X6213, and X6211 populations are presented in Supplementary Figs. S7-S10. The X6135 population was genotyped with 5,171 SNP markers, showing a dense and uniform distribution across all 21 chromosomes, averaging approximately 6 markers per Mb, with particularly good coverage on chromosomes 1D and 3D (Supplementary Fig. S7). The X6129 population contained 4,176 SNP markers, with an average density of 5 markers per Mb across the genome, in addition to 485 unmapped markers (Supplementary Fig. S8). Despite having fewer markers overall, the X6213 (2,260 markers) and X6211 (1,549 markers) populations still provided robust coverage on chromosomes 1D and 3D, enabling reliable detection of segregating loci within these genomic regions (Supplementary Figs. S9 and S10).

### Identification and Characterization of QTL for *Zymoseptoria* Resistance

#### *Stb23* on Chromosome 1DS

The first MSR locus, *Stb23* was identified on chromosome 1DS, with a peak position at 9.8 Mb (Table 2, Fig. 4a). This gene exhibited a broad impact on STB resistance, being detected for a substantial number of traits in both glasshouse and field conditions. In the X6135 population, *Stb23* was associated with 11 datasets detected under glasshouse conditions and 4 BLUPs detected in the field. For the X6213 population, seven datasets were detected in the field. The LOD scores for *Stb23* reached notable peaks, particularly in the X6135 population, where values exceeded 9 (Fig. 4a). For X6135, the phenotypic variation explained by *Stb23* ranged from 6% to 36% for the traits Nec%, Pyc%, and STB_S in the glasshouse, and 2% to 16% for the traits STB% and Pyc% in the field. For the X6213 population, *Stb23* accounted for 6% to 24% of the phenotypic variation for the traits STB% and Pyc% in the field.

**Fig. 4.**
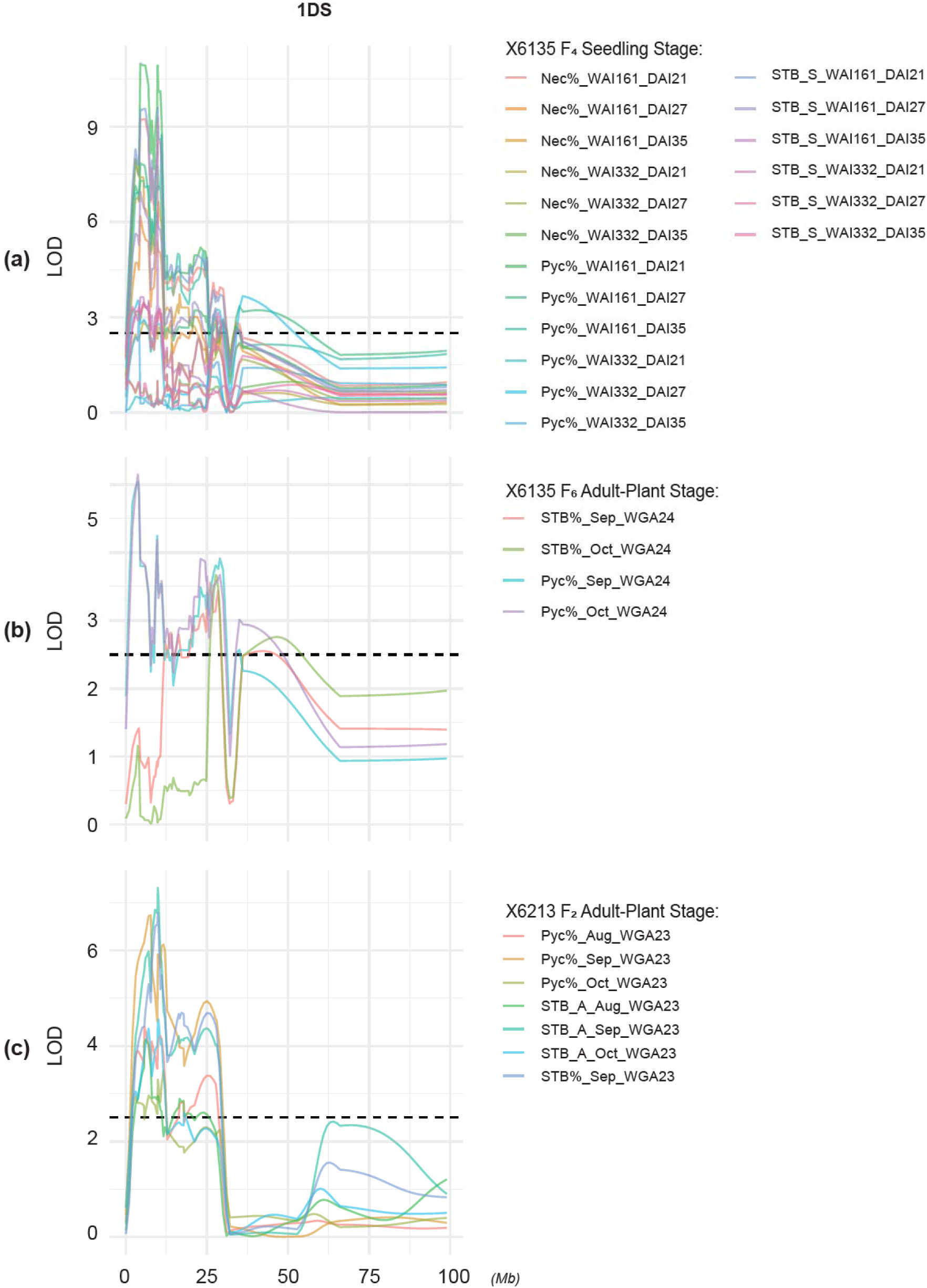
QTL mapping of the major gene *Stb23* according to IWGSC CS RefSeq v2.1 chromosome 1DS. A distance ruler indicated the position of the markers is placed at the bottom. The LOD scores for different markers were shown on the left of the figures for X6135 F_4_ population **(a)**, X6135 F_6_ population **(b)** and X6213 F_2_ **(c)**. Traits at the seedling stage include the percentage of necrotic leaf area (Nec%) on the infected leaves, the pycnidia density (Pyc%) in the necrotic leaf area, and the STB_S Scale 1 to 5. Traits were measured 21, 27, 35 days after inoculation, using two STB isolates WAI161 and WAI332. Traits at the adult-plant stage include relative maturity (Zadoks scale), STB_A scale 1-9, and the percentage of STB infected leaf area on the whole plant (STB%). Traits from adult-plant stage were measured at Wagga Wagga NSW (WGA) in 2023 and 2024.

The genetic effect of *Stb23* was characterized as semi-dominant in X6135F_4_ and X6213F_2_ populations (Table 2, Fig. 6). For a semi-dominant effect, one would expect the heterozygote to be intermediate and significantly different from both homozygotes (or one homozygote if dominance is partial). For instance, phenotypic traits Nec%, Pyc%, and STB_S in the X6135 F_4_ population, alongside STB_A in the X6213 F_2_ population (Fig. 6), exhibited significant differences between resistant (C:C) and susceptible (G:G) homozygotes (*p-value* < 0.0009). In contrast, differences between heterozygotes (G:C) and either homozygous group were substantially smaller, with 7 out of 12 genotype comparisons failing to reach significance (*p-value* > 0.05). This intermediate pattern strongly suggests a semi-dominant gene effect.

#### *Stb24* on Chromosome 3DL

The second MSR locus, *Stb24*, was mapped to chromosome 3DL, with a peak position at 512 Mb (Table 2). This gene also demonstrated significant contributions to STB resistance across multiple traits and environments. In the X6129 population, *Stb24* was detected for 6 datasets/BLUPs in both glasshouse and field conditions. Similarly, in the X6211 population, 7 datasets were detected in the field (Table 2). The LOD scores for *Stb24* showed prominent peaks, with values reaching approximately 10 in the X6129 populations (Fig. 5a). For the X6129 population, the phenotypic variation explained by *Stb24* ranged from 11% to 30% for the traits Nec%, Pyc%, and STB_S in the glasshouse and 9% to 23% for the traits Pyc% and STB% in the field. For the X6211F_2_ population, *Stb24* explained 11% to 27% of the phenotypic variation for the traits Pyc% and STB% in the field.

**Fig. 5.**
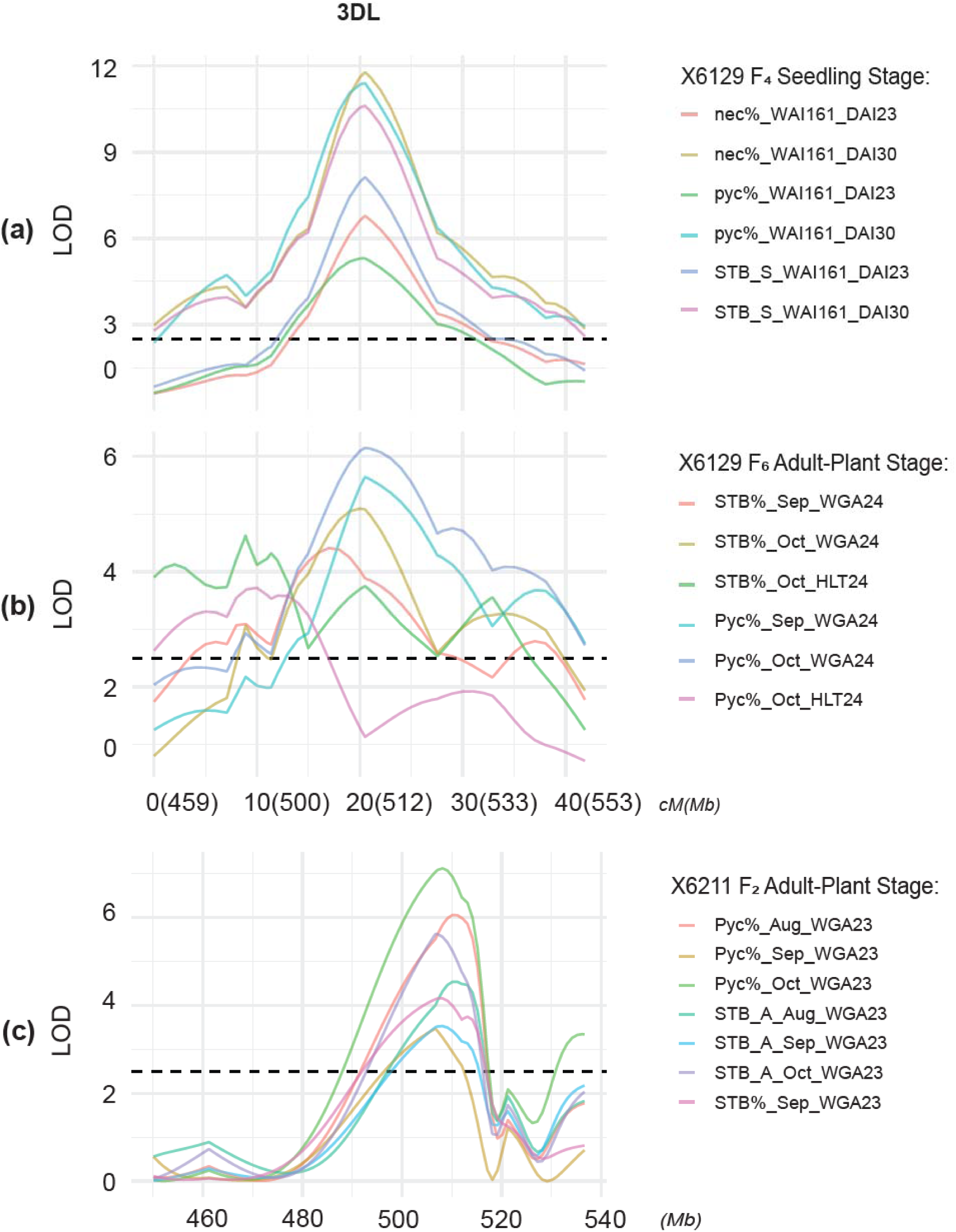
QTL mapping of the major gene *Stb24* according to IWGSC CS RefSeq v2.1 chromosome 3DL. A distance ruler indicated the position of the markers is placed at the bottom. Genetic map was constructed for the mapping of *Stb24* using X6129F_4_ population **(a)** and X6129 F_6_ population **(b)**. IWGSC CS RefSeq v2.1 physical map was used for the mapping of *Stb24* in X6211 F_2_ population **(c)**. The LOD scores for different markers were shown on the left of the figures for **(a)**, **(b)** and **(c)**. Traits at the seedling stage include the percentage of necrotic leaf area (Nec%) on the infected leaves, the pycnidia density (Pyc%) in the necrotic leaf area, and the STB_S Scale 1 to 5. Traits were measured 23 and 30 days after inoculation, using the STB isolate WAI161. Traits at the adult-plant stage include relative maturity (Zadoks scale), STB_A scale 1-9, and the percentage of STB infected leaf area on the whole plant (STB%). Traits from adult-plant stage were measured at Wagga Wagga NSW (WGA) in 2023 and 2024.

The genetic effect of *Stb24* was identified as completely dominant in the X6211 F_2_ population (Table 2, Fig. 7), confirming the dominance of the G allele. Heterozygotes (G:C) exhibited phenotypic performance similar to R homozygotes (G:G), with both groups differing significantly from S homozygotes (C:C). Specifically, no significant difference was observed between heterozygotes and R homozygotes for the STB_A trait (*p-value* > 0.059), whereas a highly significant difference was detected between heterozygotes and S homozygotes (*p-value* < 0.003)

### KASP Marker Development and Validation for *Stb23*

A KASP marker, designated snp*_*1D1217527 which is tightly linked to *Stb23* (Table 3), was developed for the evaluation of additive gene effects (Fig. 6). The polymorphism detected by snp*_*1D1217527 was confirmed using both the X6135 F₄ segregating population and the AuSTB panel (Fig. 6a). Following inoculation with isolate WAI161, significant differences were observed between the resistant (C:C) and susceptible (G:G) classes for Nec%, Pyc%, and STB_S. For Nec%_WAI161_DAI21, the *p-value* was 5.93×10⁻⁹; for Pyc%_WAI161_DAI21, 1.33×10⁻¹²; and for STB_S_WAI161_DAI21, 7.20×10⁻⁹ (Fig. 6b). Additional analyses across multiple time points and using isolates WAI161 and WAI332 (Supplementary Figs. S13 and S14) consistently supported the strong gene effects. For example, Nec%_WAI161_DAI35 showed a *p-value* of 2.38×10⁻⁶ (Supplementary Fig. S13), and Pyc%_WAI332_DAI21 showed a *p-value* of 2.35×10⁻⁴ (Supplementary Fig. S14). Validation was next extended to the adult-plant stage using the X6135 F₆ population (Fig. 6c). Comparisons of additive gene effects between the C:C and G:G groups revealed significant differences for Pyc%_Sep_WGA24 (*p-value* = 6.83×10⁻⁶) and Pyc%_Oct_WGA24 (*p-value* = 8.28×10⁻⁶).

**Table 3.**
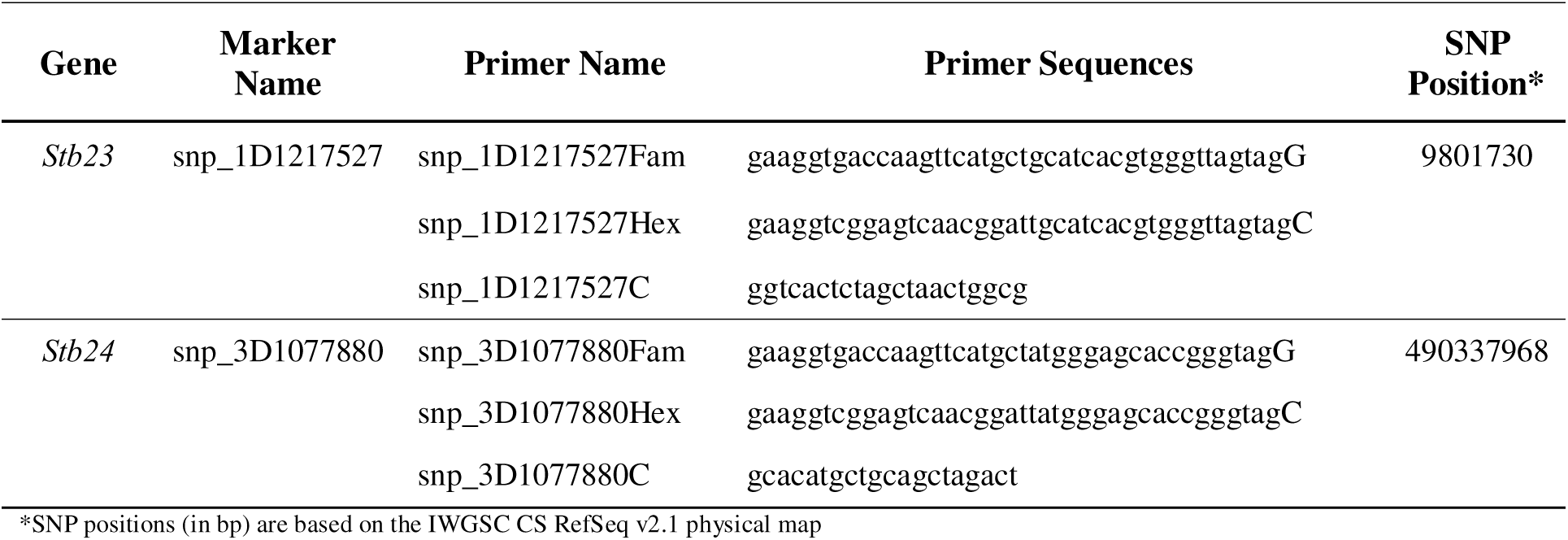
KASP marker primers.

**Fig. 6.**
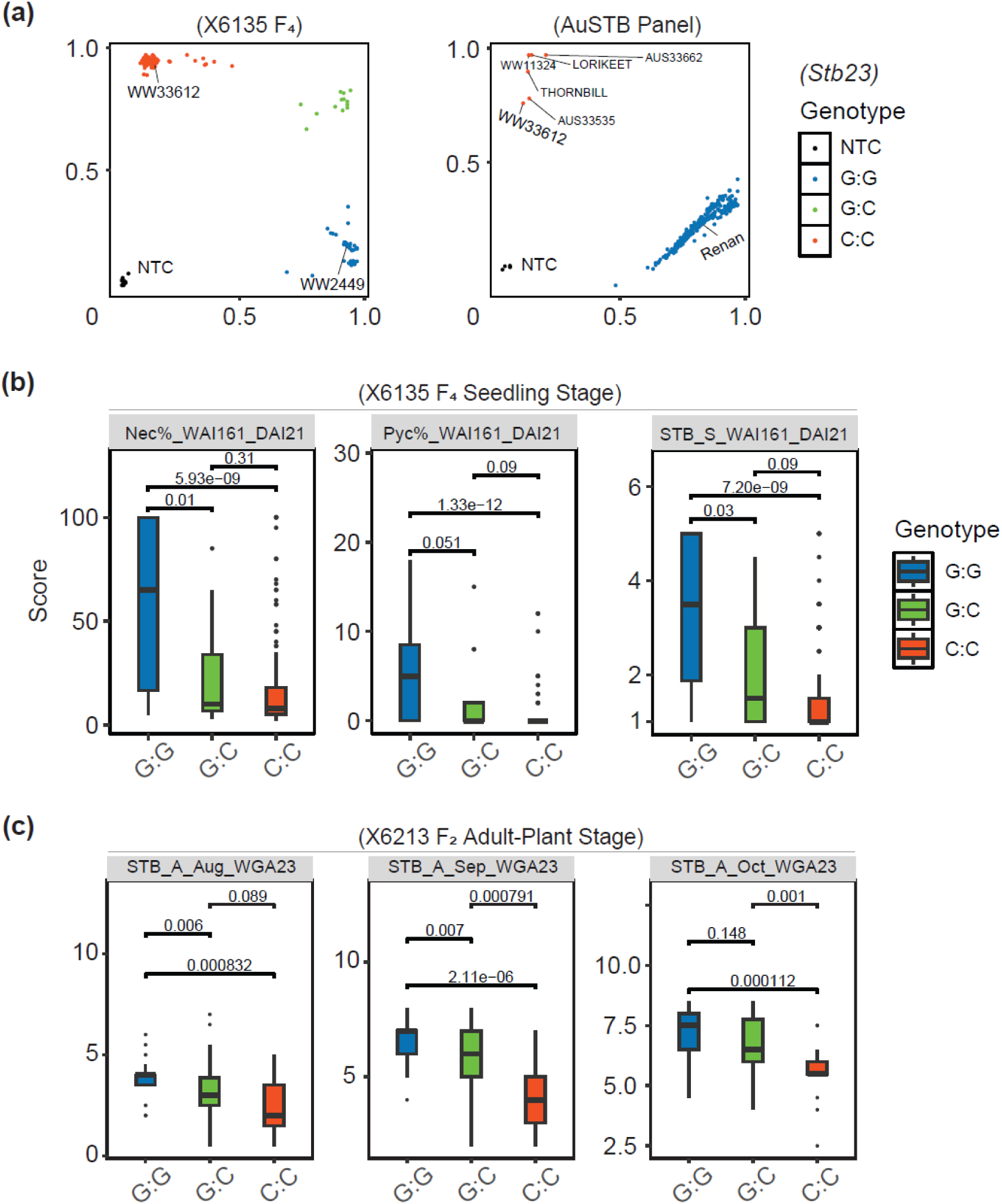
Validation of the KASP marker snp_1D1217527 tightly linked to *Stb23* in segregating populations and AuSTB panel. **(a)** snp_1D1217527 was genotyped in X6135 F_4_ segregating population and AuSTB panel to validate the polymorphism. **(b)** Comparisons of additive effects among homozygous resistant, susceptible and heterozygous lines in X6135 F_4_ population at the seedling stage. **(c)** Comparisons of additive effects among homozygous resistant, susceptible and heterozygous lines in X6213 F_2_ population at the adult-plant stage. Traits from adult-plant stage were measured at Wagga Wagga NSW (WGA) in 2023 and 2024. Red color represents homozygous resistant lines (C:C), green color represents the heterozygous lines (G:C), and blue color represents the homozygous susceptible lines (G:G). The *p* values acquired from multiple Wilcoxon test are shown above each pair of homozygous resistance and susceptible lines in each family.

A second, independent validation further confirmed that *Stb23* significantly contributes to disease reduction, reinforcing its role as a major MSR locus. This validation employed BC₁F₂ homozygous lines derived from three distinct genetic backgrounds: X6284-6, X6312-10, and X6312-2 (Table 1). As shown in Fig. 8a, comparisons of resistant (C:C) versus susceptible (G:G) homozygotes consistently demonstrated significant additive effects for Pyc% and STB% in the 2024 field trials at Wagga Wagga (WGA). In the X6284-6 family, Pyc%_Sep_WGA24 displayed a *p-value* of 5×10⁻³ (9 resistant vs 6 susceptible). In X6312-10, Pyc%_WGA24 showed a *p-value* of 9×10⁻³ (13 resistant vs 12 susceptible), and in X6312-2 the *p-value* was 0.001 (16 resistant vs 13 susceptible). Supplementary Fig. S19a provides additional supporting data, including *p-value*s of 0.073 for Pyc%_Sep_WGA24 and 2.24×10⁻⁴ for STB%_Sep_WGA24 in X6284-6.

Together, these results demonstrate a strong, consistent, and reproducible additive effect of *Stb23* across multiple developmental stages and diverse genetic backgrounds, confirming its stability and value as a robust multi-stage resistance (MSR) locus.

### KASP Marker Development and Validation for *Stb24*

A KASP marker, snp_3D1077880 which is tightly linked to *Stb24* (Table 3), was developed for the gene effects (Fig. 7). The allelic polymorphism detected by this marker was confirmed by genotyping the X6129 F₄ segregating population and the AuSTB panel (Fig. 7a). Following inoculation with isolate WAI161, resistant (G:G) and susceptible (C:C) homozygous groups showed highly significant differences for Nec%, Pyc%, and STB_S. For Nec%_WAI161_DAI28, the *p-value* was 9.11×10⁻⁹; for Pyc%_WAI161_DAI28, 1.15×10⁻¹⁰; and for STB_S_WAI161_DAI28, 1.66×10⁻⁹. Adult-plant validation using the X6129 F₆ population (Fig. 7c) further confirmed the contribution of *Stb24* to field resistance. Significant additive gene effects were detected for Pyc%_Oct_HLT24 (*p-value* = 0.076), Pyc%_Oct_WGA24 (*p-value* = 1.63×10⁻⁵), and Pyc%_Sep_WGA24 (*p-value* = 0.001). Additional support was obtained from the X6211 F₂ population, which showed consistent associations across Pyc%, STB_A, and STB% traits, including a *p-value* of 3.01×10⁻⁵ for Pyc%_Aug_WGA23 (Supplementary Fig. S18).

**Fig. 7.**
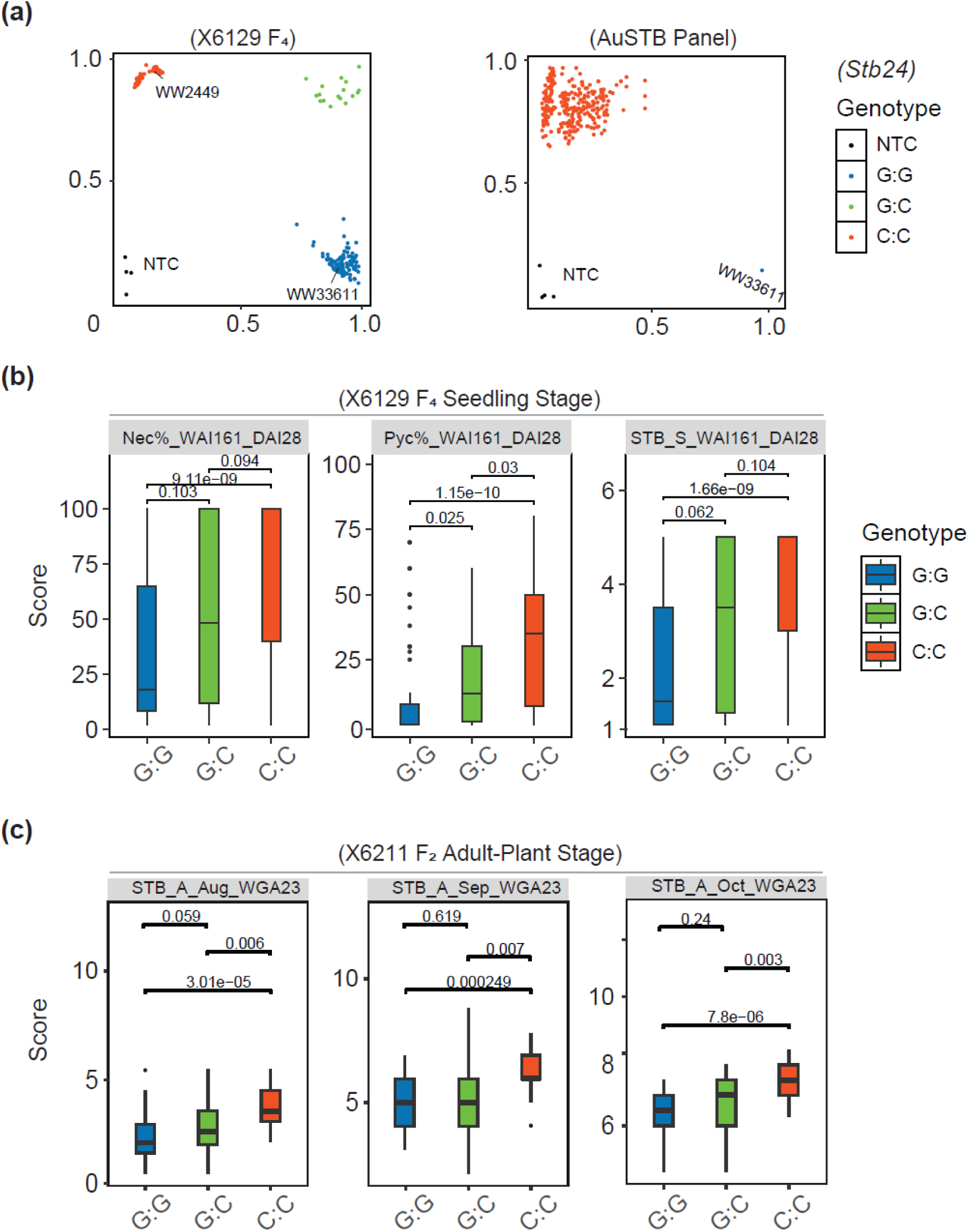
Validation of the KASP marker snp_3D1077880 tightly linked to *Stb24* in segregating populations and AuSTB panel. **(a)** snp_3D1077880 was genotyped in X6129 F_4_ segregating population and AuSTB panel to validate the polymorphism. **(b)** Comparisons of additive effects among homozygous resistant, susceptible and heterozygous lines in X6129 F_4_ population at the seedling stage. **(c)** Comparisons of additive effects among homozygous resistant, susceptible and heterozygous lines in X6211 F_2_ population at the adult-plant stage. Traits from adult-plant stage were measured at Wagga Wagga NSW (WGA) in 2023 and 2024. Blue color represents homozygous resistant lines (G:G), green color represents the heterozygous lines (G:C), and red color represents the homozygous susceptible lines (C:C). The *p* values acquired from multiple Wilcoxon test are shown above each pair of homozygous resistance and susceptible lines in each family.

A second validation step further substantiated the role of *Stb24* using BC₁F₂ homozygous plants derived from four distinct genetic backgrounds: X6285-3, X6285-5, X6293-5, and X6300-1 (Table 1). As shown in Fig. 8b, resistant (G:G) and susceptible (C:C) plants displayed clear differences in Pyc% and STB% during the 2024 field assessments at Wagga Wagga (WGA), although the additive gene effects were weaker than *Stb23*. For example, in the X6285-3 family, Pyc%_WGA24 showed a *p-value* of 0.0522 (9 resistant vs 6 susceptible), while X6285-5 exhibited a *p-value* of 0.0554 (26 resistant vs12 susceptible), and X6293-5 showed a *p-value* of 0.0244 (15 resistant vs 12 susceptible).

**Fig. 8.**
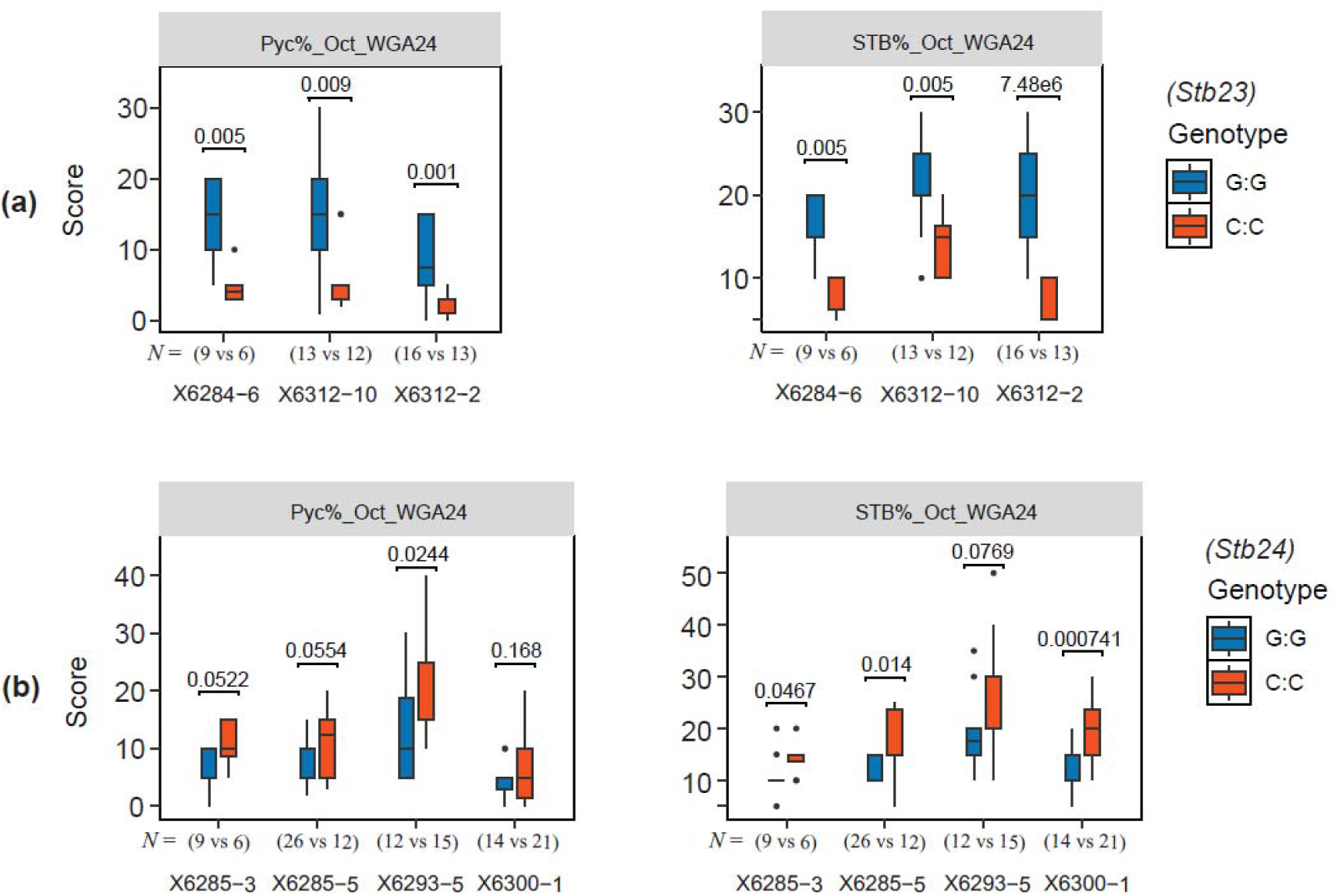
Comparisons of additive effects between homozygous resistant lines and homozygous susceptible lines on *Stb23* and *Stb24*. **(a)** Comparisons of additive effects of *Stb23* among three different F_2_ families X6284-6, X6312-10, and X6312-2. Red color represents homozygous resistant lines (C:C) and blue color represents the homozygous susceptible lines (G:G). **(b)** Comparisons of additive effects of *Stb24* among four different F_2_ families, X6285-3, X6285-5, X6293-5, and X6300-1. Blue color represents homozygous resistant lines (G:G) and red color represents the homozygous susceptible lines (C:C). The number of homozygous lines used in the analysis shown under the bars. The *p* values acquired from multiple Wilcoxon test are shown above each pair of homozygous resistance and susceptible lines in each family.

Collectively, these results demonstrate a strong, consistent, and reproducible additive gene effect of *Stb24* across developmental stages and across diverse genetic backgrounds, confirming the stability and utility of this locus as a robust multi-stage resistance (MSR) gene.

### Mapping and Annotation of *Stb23* and *Stb24* Candidates

For the *Stb23* locus on chromosome 1DS, 24 candidates from a total 185 genes were identified from within the QTL region (Supplementary Table S4). These genes include a variety of resistance-related gene families, such as serine/threonine-protein kinases, rust resistance kinases (e.g., Lr10-like), LRR receptor-like kinases, and NBS-LRR-like proteins. Notably, two wall-associated receptor kinase 2-like (WAKL) genes similar to *Stb6/TaWAKL4* (Saintenac et al. 2018) and four G-type lectin S-receptor-like kinases similar to *Stb15* (Hafeez et al. 2025) were present, suggesting their potential roles in host defense responses.

Similarly, for the *Stb24* locus on chromosome 3DL, 10 candidates out of a total of 125 genes were identified within the QTL region (Supplementary Table S4). The descriptions of these genes primarily include various types of LRR-like protein kinases, and serine/threonine-protein kinases, as well as an MLO-like protein. This set of genes suggests novel or less-understood resistance mechanisms within this region. Further investigation into these candidate genes will be crucial for elucidating the molecular basis of *Stb24*-mediated resistance.

## Discussion

This study successfully identified and validated two significant resistance loci, one on chromosome 1DS (designated *Stb23*) and one on chromosome 3DL (designated *Stb24*). Both loci confer stable MSR to *Zymoseptoria tritici* across multiple growth stages (Figs. 6a and 7a). The development and validation of two tightly linked KASP markers, snp_1D1217527 for *Stb23* (Fig. 6) and snp_3D1077880 for *Stb24* (Fig. 7), represent a substantial advancement for marker-assisted breeding (MAB) for STB resistance, consistent with the increasing emphasis on efficient and reliable MAB strategies.

The *Stb23* locus on chromosome 1DS is positioned at 9.8 Mb and originates from the synthetic wheat line ‘WW33612’, derived from ‘W7984’ (Table 2; Fig. 6a). Our study provides a robust characterisation of this important resistance locus, demonstrating its significant contribution to resistance across multiple populations and environments (Fig. 6a). The locus exhibits stable effects across seven populations in various field and greenhouse environments, and this work delivers the first validated KASP marker enabling its deployment in wheat breeding. By converting a previously tentative genetic signal into a reliable and selectable allele, we substantially enhance the practical utility of this locus.

The *Stb23* region co-localises with several previously reported QTLs, most notably QStb.ipk-1D, which also originates from ‘W7984’ (Simón et al. 2004). However, QStb.ipk-1D was originally characterised by modest LOD scores (approximately 3) and a broad, unresolved interval that lacked subsequent validation. This region has also been associated with three additional loci: QStb.B22-1D.a (Naz et al. 2015), Qstb.renan-1D (Langlands-Perry et al. 2022), and *Stb19* (Yang et al. 2018). QStb.B22-1D.a, with resistance derived from the synthetic accession ‘Syn022L’, was reported to explain 15.1% of phenotypic variance in field trials (Naz et al. 2015). Although QStb.B22-1D.a could be allelic to *Stb23*, confirmation is not currently feasible because its tightly linked marker, GENE_0014_822, is not publicly available.

Qstb.renan-1D is also located within the 7-11 Mb interval on the IWGSC CS RefSeq v2.1 physical map (Langlands-Perry et al. 2022). Given that ‘Renan’ carries a well-documented *Aegilops ventricosa* translocation harbouring multiple resistance genes (Robert et al. 1999), allelism between Qstb.renan-1D and *Stb23* remains plausible.

The KASP marker snp_1D1217527 developed in this study does not differentiate the resistant allele in ‘WW33612’ from those in ‘WW11324’, ‘Thornbill’, or ‘AUS33535’, nor does it distinguish *Stb23* from the *Stb19*-carrying cultivars ‘Lorikeet’ and ‘AUS33662’. However, *Stb23* is confirmed to be distinct from *Stb19*, as a separate marker (snp_1218021) (Yang et al. 2018) is able to discriminate between *Stb19* and *Stb23* (data not shown). In addition, the different reactions to WAI161 isolate test at the seedling stage further indicate they are different genes. Recent fine-mapping efforts within the chromosome 1DS region suggest a high likelihood that ‘WW33612’ and ‘AUS33535’ both carry the same resistance gene *Stb23*. This inference is further supported by their highly comparable field performance in independently derived BC₂F₂ lines from the respective resistant parents (data not shown). Further resolution of this region will require fine mapping and targeted genome sequencing to clarify the gene content within the 1DS interval across these synthetic wheat backgrounds.

The locus on chromosome 3DL, designated *Stb24*, is positioned between 506-516 Mb and originates from the line ‘WW33611’, which traces back to ‘Fleche D’or’ (Table 2; Fig. 7a). A nearby QTL, QStb.ipk-3D, with resistance contributed by ‘W7984’, has been reported within the 520-605 Mb region of chromosome 3D on the CS RefSeq v2.1 physical map. This QTL was detected using three traits with LOD scores below 3, based on a single isolate (IPO 92067) assessed at two developmental stages (Simón et al. 2004). As with QStb.ipk-1D, the absence of subsequent validation prevents definitive assessment of whether QStb.ipk-3D is allelic to *Stb24*. However, evidence from this study effectively rules out this possibility: since *Stb23* was identified from ‘W7984’ on chromosome 1DS, no corresponding resistance was detected on 3DL from this donor background in our experiments (data not shown).

The Meta-QTL MQTL15, which spans 480-570 Mb on chromosome 3D, overlaps broadly with the *Stb24* interval. However, its large physical range and limited resolution impede precise comparison with our finely mapped locus. This study provides the first robust characterisation of *Stb24*, supported by high LOD values (∼10), consistent phenotypic effects across multiple environments, and the development of the first validated KASP marker for this genomic region-transforming a previously diffuse genetic signal into a reliable and readily deployable breeding tool. Furthermore, BLAST-based searches of candidate genes in the *Stb24* interval revealed no WAKL genes resembling *Stb6*/*TaWAKL4* (Saintenac et al. 2018) nor any CRK-like genes similar to *Stb16q*/*TaCRK6* (Saintenac et al. 2021) (Supplementary Table S4). This suggests that *Stb24* may represent a novel form of resistance gene, distinct from all *Stb* genes cloned to date.

Pycnidia density on infected leaves (Pyc%) is a robust and highly informative trait for genetic analysis, often producing stronger QTL signals and higher LOD scores than necrotic leaf area (Nec%) or whole-plant infection severity (STB%) (Figs. 6 and 7). This pattern is consistent with previous studies, where 8 of 13 MSR-associated QTLs were linked to pycnidia sporulation (Yang et al. 2022), and with Thauvin et al. (2024), who reported that 20 of 40 QTLs were detected for both necrosis and sporulation using the same isolate. Biologically, *Z. tritici* isolates resulting in higher pycnidia densities are expected to generate more asexual spores, potentially causing greater cumulative damage over successive infection cycles (Stewart et al. 2016). Given the strong correlations observed among Pyc%, Nec%, and STB% in this and other genetic studies (Thauvin et al. 2024; Yang et al. 2022; Yates et al. 2019), QTLs detected using pycnidia-related traits are reliable predictors of broader resistance effectiveness.

For *Z. tritici*, resistance expressed at the seedling stage frequently fails to translate into effective protection at the adult-plant stage under field conditions. Thauvin et al. (2024) demonstrated that only one of twenty seedling-stage QTL identified using isolates IPO09415 and INRA16-TM0229 was subsequently detected in field trials. Similarly, our previous GWAS analysis showed that only four of thirteen seedling-detected loci were retained in adult-plant evaluations (Yang et al. 2022). To address this persistent translational gap and increase the likelihood of identifying MSR or APR loci with genuine breeding value, we employed a paired-population strategy within a semi-double-blind framework. In this design, BC₁F₂ populations derived from resistant F₂ individuals were evaluated for field-based resistance, while their corresponding F₄ descendants were phenotyped for seedling resistance without prior knowledge of the underlying loci (Fig. 1). This coordinated, stage-specific evaluation substantially reduces the risk of prioritising loci that act only at the seedling stage and therefore have limited utility for breeding. At the same time, it accelerates the progression from locus discovery to validation and enables rapid deployment of newly identified *Stb* MSR genes into breeding pipelines. Notably, this practice can reduce the timeframe required to deliver validated resistance traits into advanced breeding materials from approximately seven years to three years.

## Conclusions

This study characterises two valuable resistance loci: the gene *Stb23* on chromosome 1DS and the gene *Stb24* on chromosome 3DL. Both loci confer stable and consistent resistance to *Zymoseptoria tritici* across seedling and adult-plant growth stages. *Stb23*, located at 9.8 Mb on 1DS, exhibits additive genetic effects and explains a substantial proportion of phenotypic variation (6-36% in glasshouse assessments and 2-16% in field trials). In contrast, *Stb24*, mapped to 512 Mb on 3DL, displays dominant genetic effects, accounting for 11-30% of variation in glasshouse evaluations and 9-23% under field conditions.

A major outcome of this research is the successful development and rigorous validation of tightly linked KASP markers-snp_1D1217527 for *Stb23* and snp_3D1077880 for *Stb24*. These markers performed reliably across diverse genetic backgrounds and developmental stages, confirming the robustness of both loci. The multi-stage resistance conferred by *Stb23* and *Stb24*, combined with efficient marker tools and their genetic effects, will substantially accelerate the incorporation of these genes into elite wheat germplasm through marker-assisted breeding.

## Author contribution statement

NY and AM conceived of the study and coordinated its design. The project was led by AM. NY executed the experiments, collected and analysed the data. NY wrote the draft, with contributions from BO, BB, PS and AM. All authors read and approved the final manuscript.

## Acknowledgments

The Authors thank M McCaig for the management of field trials, T Goldthorpe, R Beecher and B McVittie for their technical contributions on the collection of leaf samples and phenotypic data. The authors also thank Dr A Kilian from DArT Pty Ltd for detailed advice on genotyping. This study was conducted as a co-investment between the NSW Department of Primary Industries Regional Development and the Grain Research and Development Corporation, under projects DPI2304-007RTX and DPI2305-008RTX of the Grains, Agronomy and Pathology Partnership.

## Declaration of generative AI in scientific writing

The authors used ChatGPT and Gemini as a tool to improve linguistic clarity during manuscript preparation. All AI-assisted text was critically assessed, edited, and validated by the authors, who accept full responsibility for the accuracy and integrity of the final article.

## Compliance with ethical standards

The authors declare that the research was conducted in the absence of any commercial or financial relationships that could be construed as a potential conflict of interest.

