## Supplementary Figures and Legends for "Two Novel Genes, *Stb23* and *Stb24*, Conferring Multi-stage Resistance to *Zymoseptoria tritici*: Rapid Deployment in Marker-Assisted Wheat Breeding"

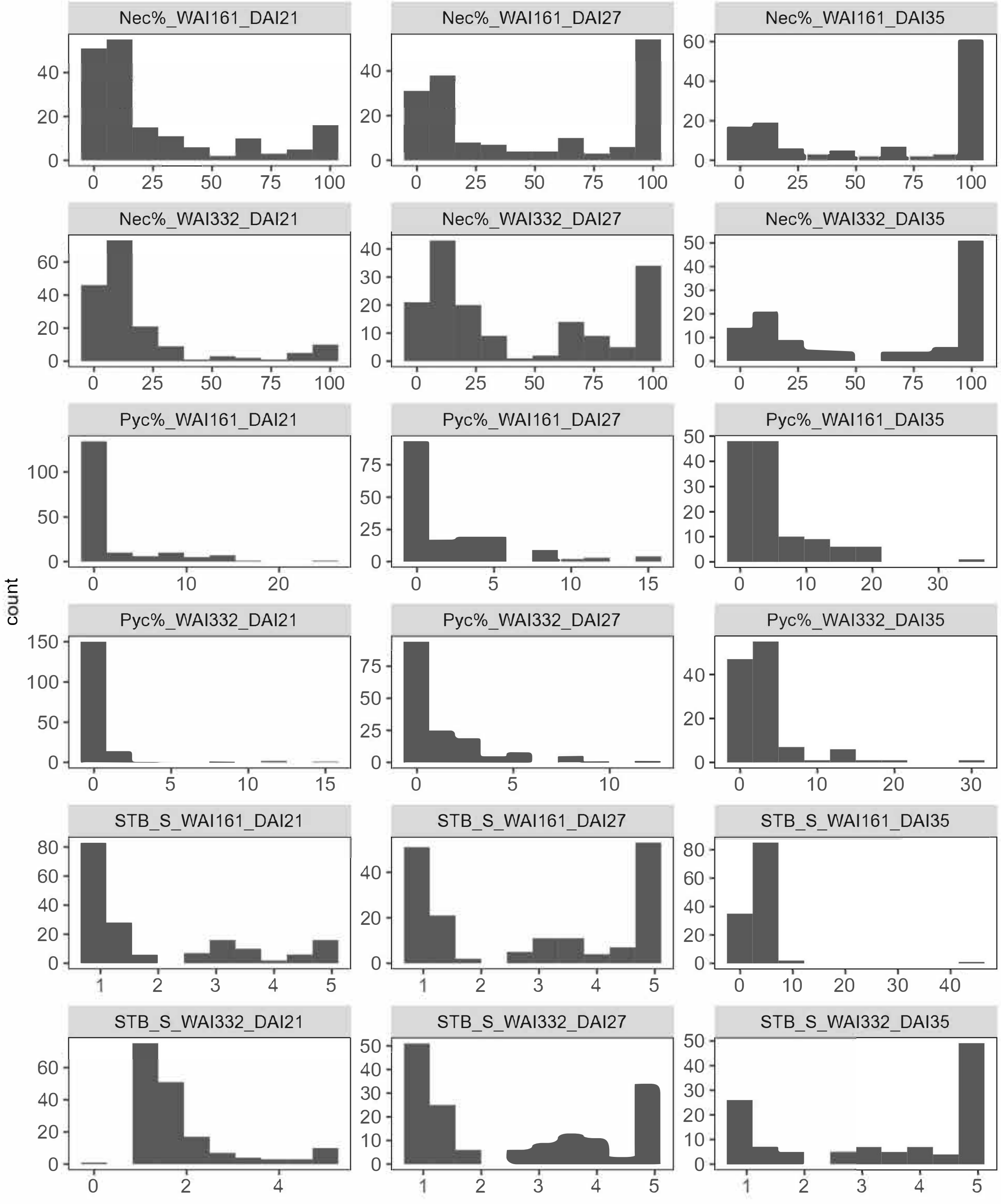


**Supplementary Fig. S1** Histogram analysis of 18 datasets on the X6135 $F_4$ segregating population. Traits at the seedling stage include the percentage of necrotic leaf area (Nec%) on the infected leaves, the pycnidia density (Pyc%) in the necrotic leaf area, and the STB_S Scale 1 to 5. Traits were measured at 21, 27 and 35 days after inoculation, using the isolate WAI161 and WAI332.

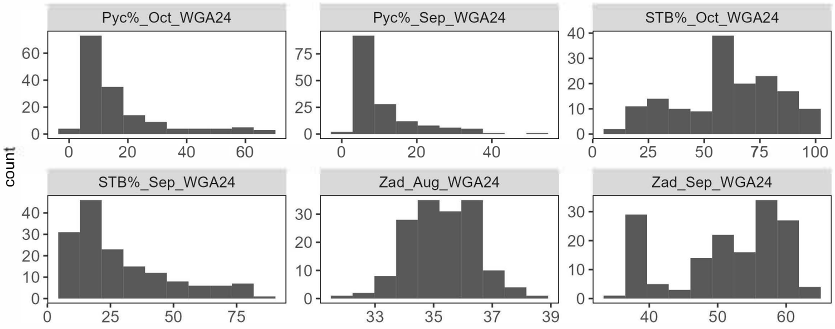


**Supplementary Fig. S2** Histogram analysis of six BLUPs on the X6135 $F_6$ segregating population. Traits at the adult-plant stage include Relative maturity (Zadoks scale) and the percentage of STB infected leaf area on the whole plant (STB%). Traits were measured at Wagga Wagga NSW (WGA) in 2024.

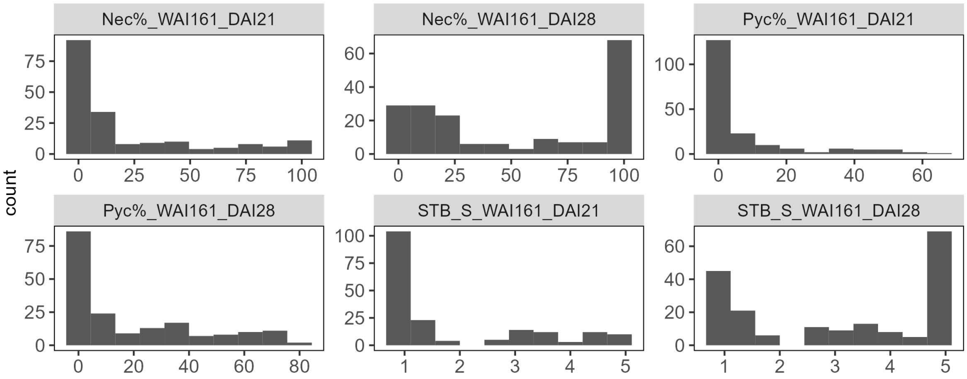


**Supplementary Fig. S3** Histogram analysis of six datasets on the X6129 $F_4$ segregating population. Traits at the seedling stage include the percentage of necrotic leaf area (Nec%) on the infected leaves, the pycnidia density (Pyc%) in the necrotic leaf area, and the STB_S Scale 1 to 5. Traits were measured at 21 and 28 days after inoculation, using the isolate WAI161.

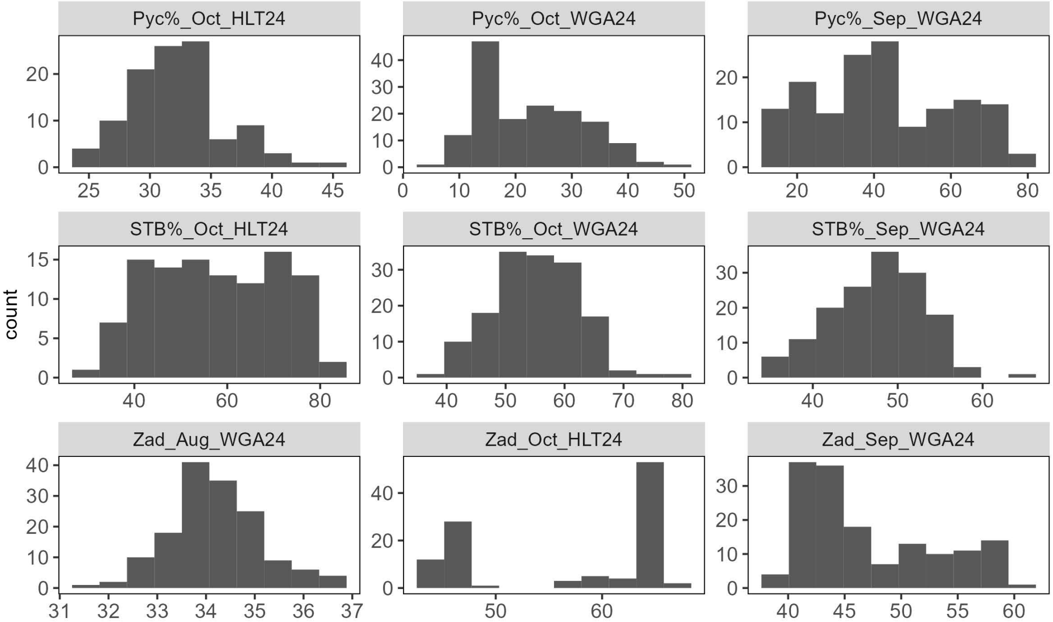


**Supplementary Fig. S4** Histogram analysis of nine BLUPs on the X6129 $F_6$ segregating population. Traits at the adult-plant stage include Relative maturity (Zadoks scale) and the percentage of STB infected leaf area on the whole plant (STB%). Traits were measured at Wagga Wagga NSW (WGA) and Hamilton Vic (HLT) in 2024.

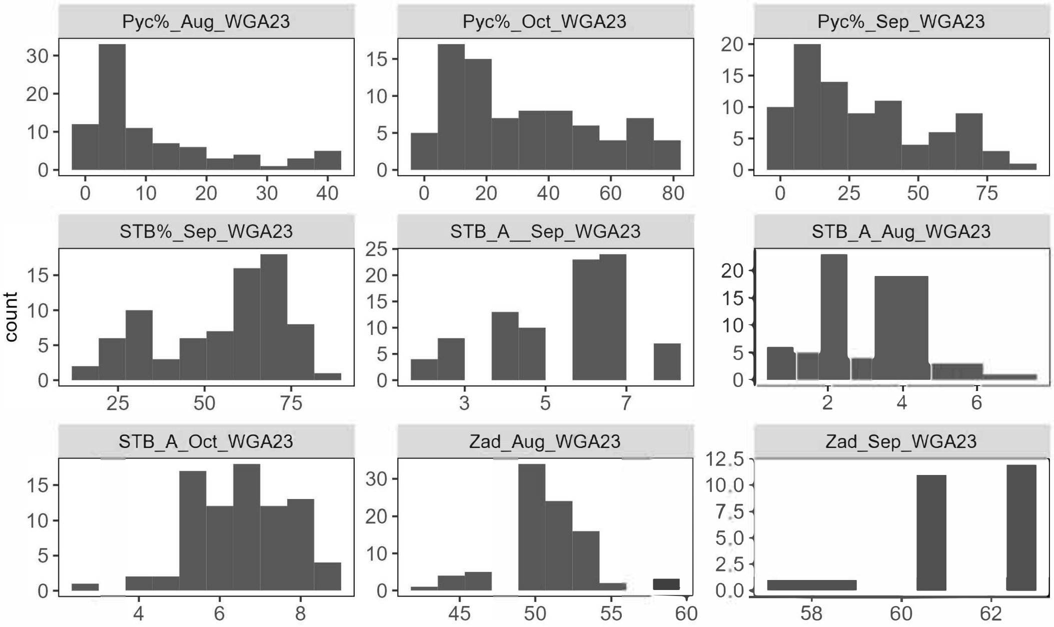


**Supplementary Fig. S5** Histogram analysis of nine datasets on the X6213 $F_2$ segregating population. Traits at the adult-plant stage include Relative maturity (Zadoks scale) and the percentage of STB infected leaf area on the whole plant (STB%). Traits were measured at Wagga Wagga NSW (WGA) in 2023.

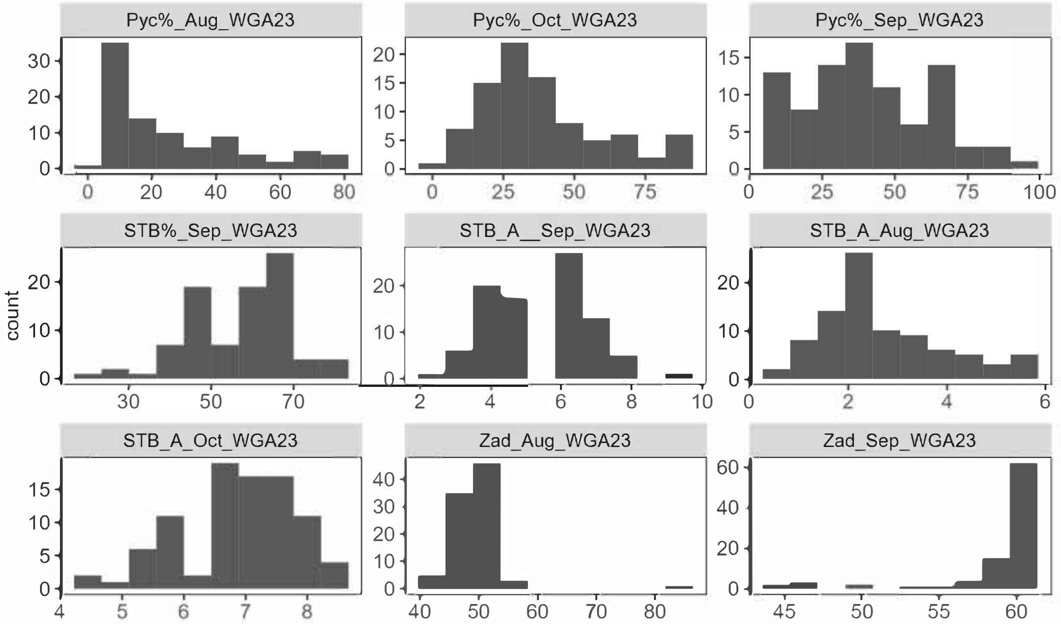


**Supplementary Fig. S6** Histogram analysis of nine datasets on the X6211 $F_2$ segregating population. Traits at the adult-plant stage include Relative maturity (Zadoks scale) and the percentage of STB infected leaf area on the whole plant (STB %). Traits were measured at Wagga Wagga NSW (WGA) in 2023.

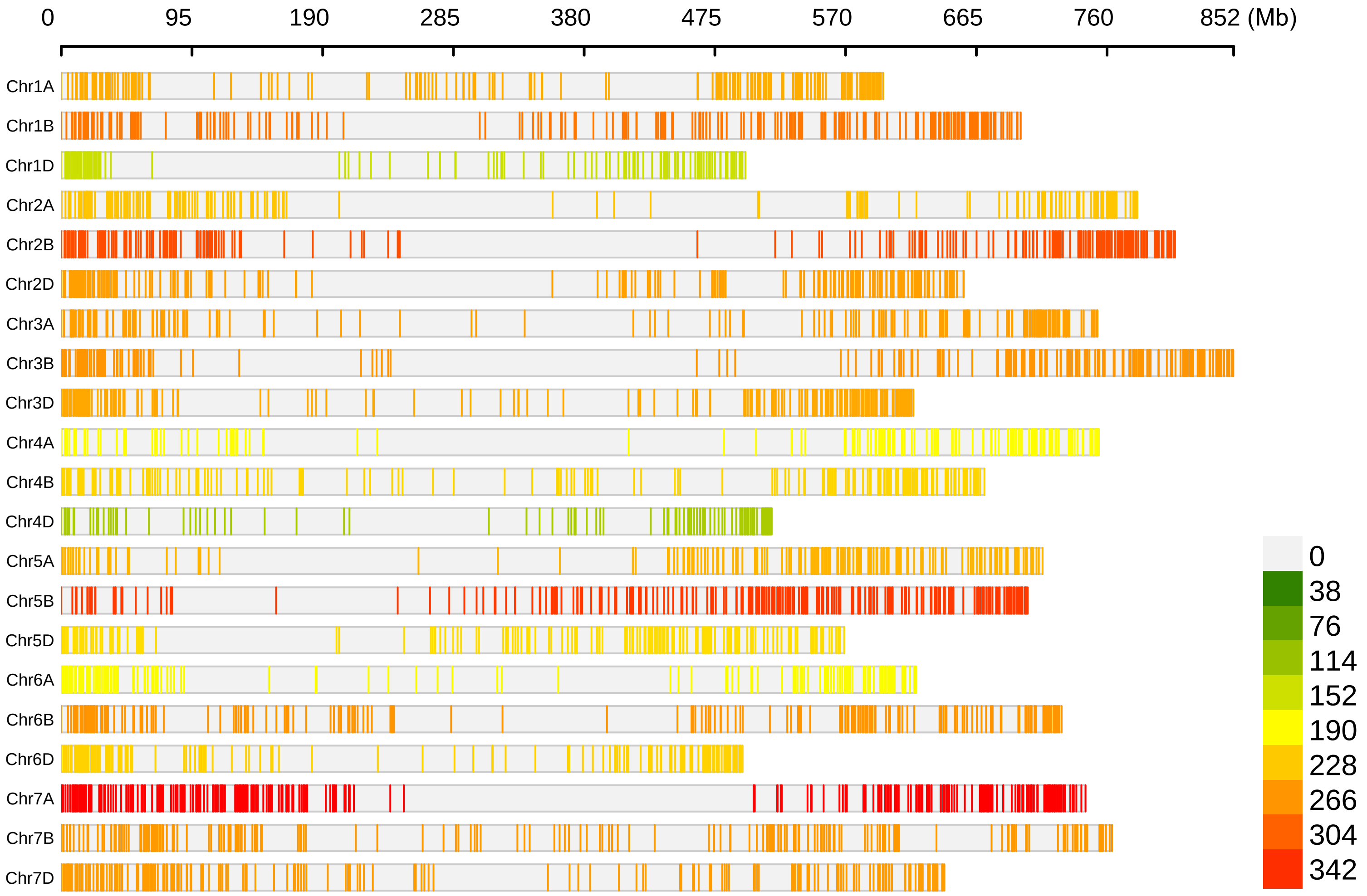


**Supplementary Fig. S7** Distribution and density of polymorphic SNP markers on 21 chromosomes for X6135 segregating population.

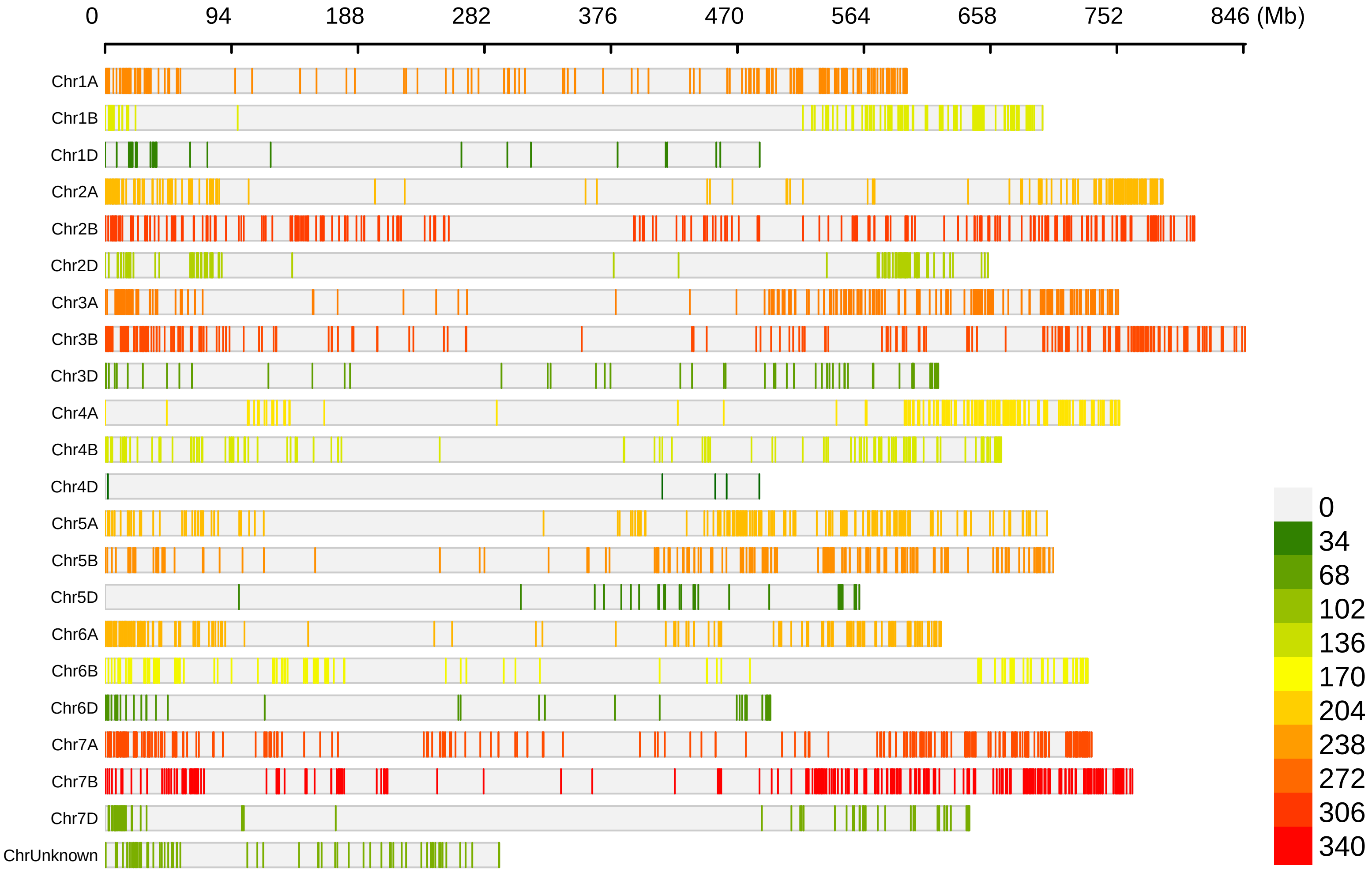


**Supplementary Fig. S8** Distribution and density of polymorphic SNP markers on 21 chromosomes and unmapped chromosome for X6129 segregating population.

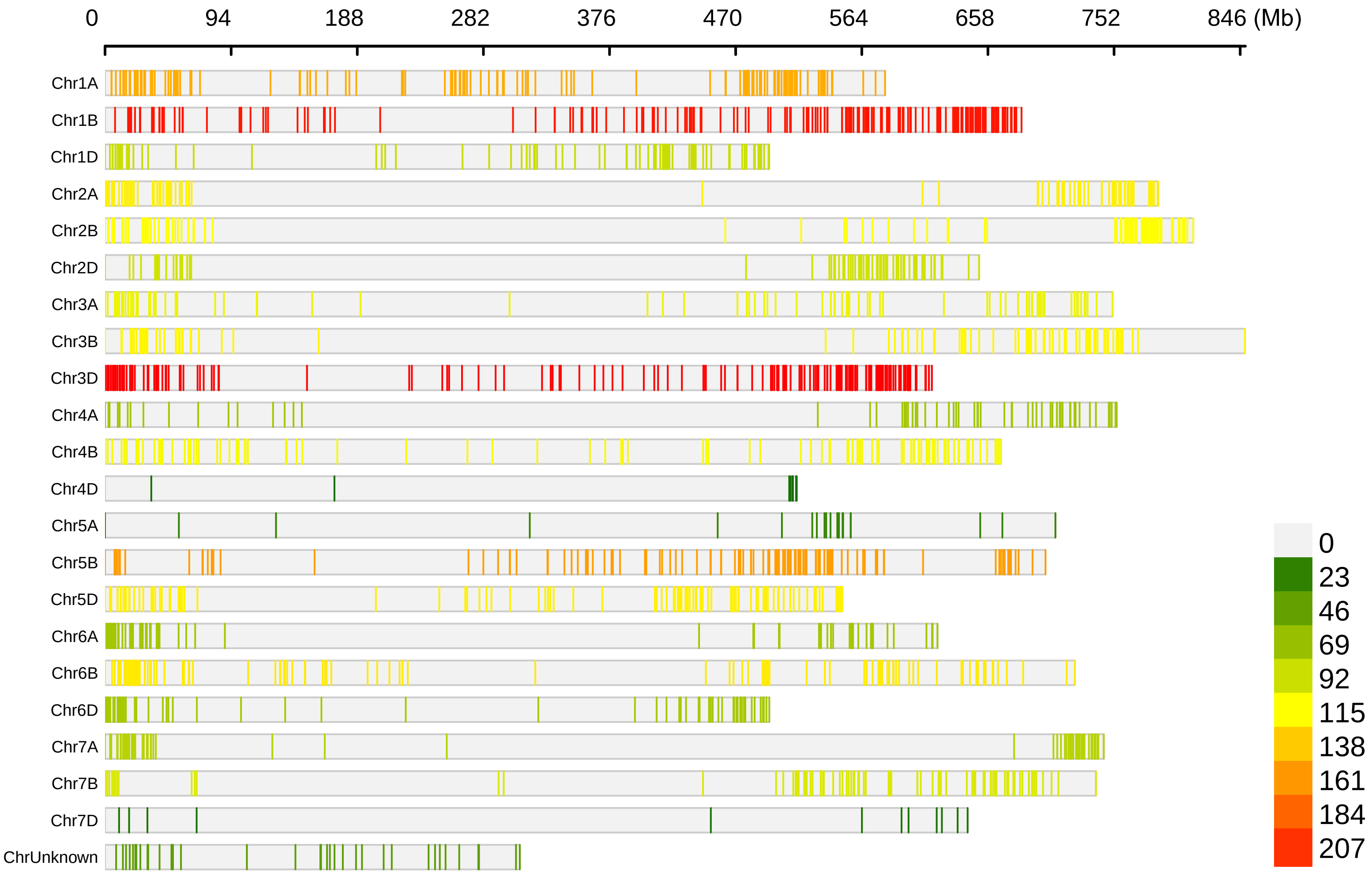


**Supplementary Fig. S9** Distribution and density of polymorphic SNP markers on 21 chromosomes and unmapped chromosome for X6213 $F_2$ segregating population.

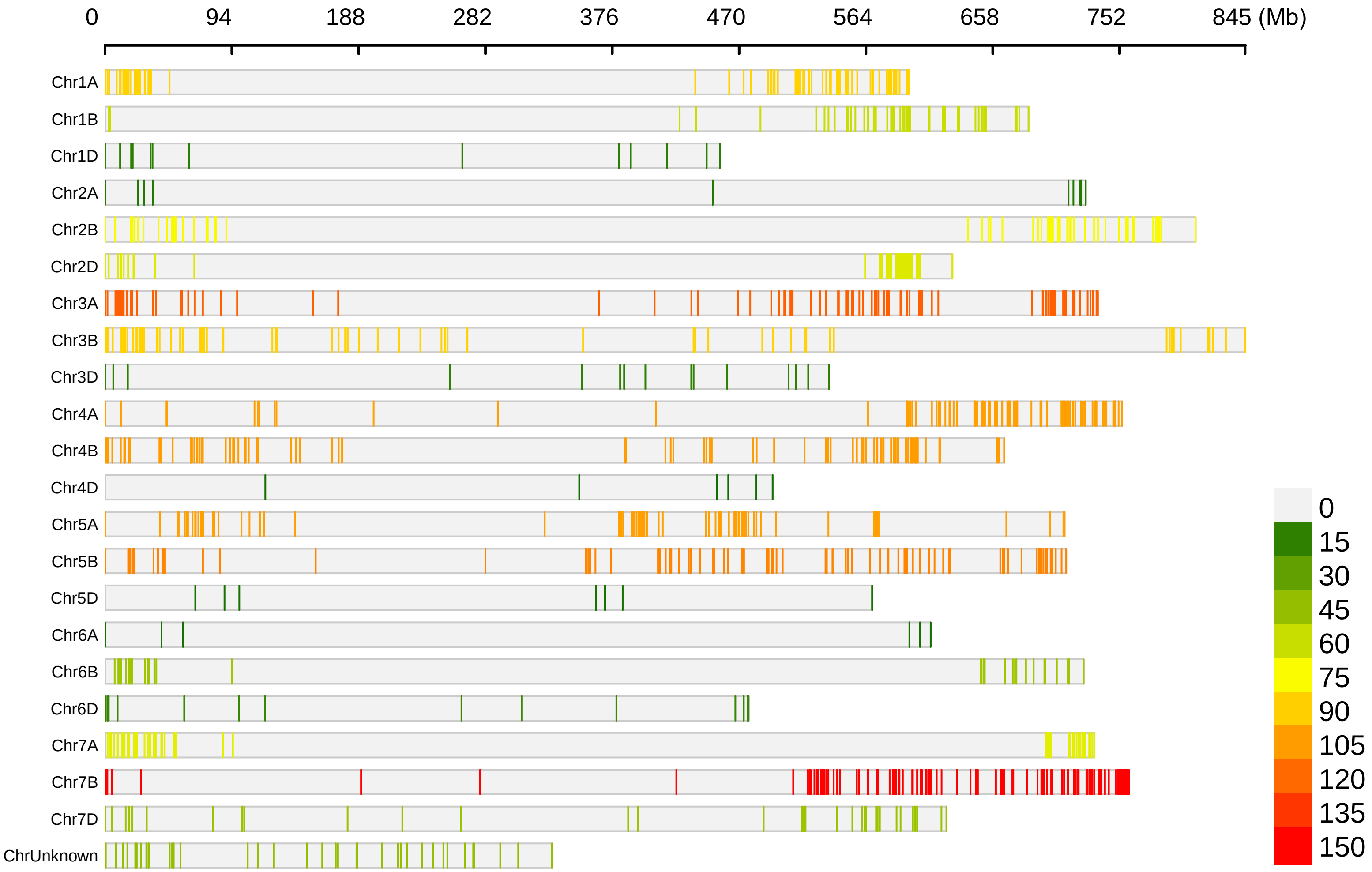


**Supplementary Fig. S10** Distribution and density of polymorphic SNP markers on 21 chromosomes and unmapped chromosome for X6211 $F_2$ segregating population

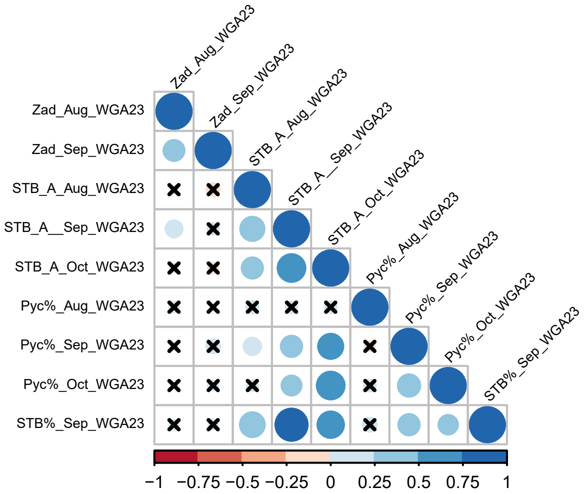


**Supplementary Fig. S11:** Correlation analysis between various datasets measured on populations X6213. Correlation analysis between 9 datasets measured at Adult-plant stage. Traits at the adult-plant stage include Relative maturity (Zadoks scale), STB_A scale 1-9, and the percentage of STB infected leaf area on the whole plant (STB%). Traits were measured at Wagga Wagga NSW (WGA) in 2023.

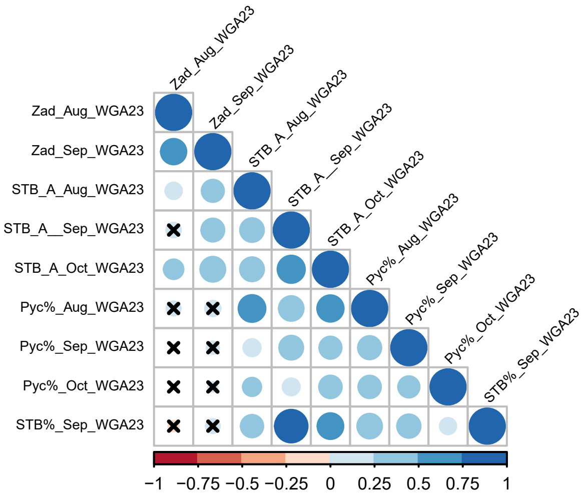


**Supplementary Fig. S12** Correlation analysis between various datasets measured on populations X6211. Correlation analysis between 9 Bdatasets measured at Adult-plant stage. Traits at the adult-plant stage include Relative maturity (Zadoks scale), STB_A scale 1-9, and the percentage of STB infected leaf area on the whole plant (STB%). Traits were measured at Wagga Wagga NSW (WGA) in 2023.

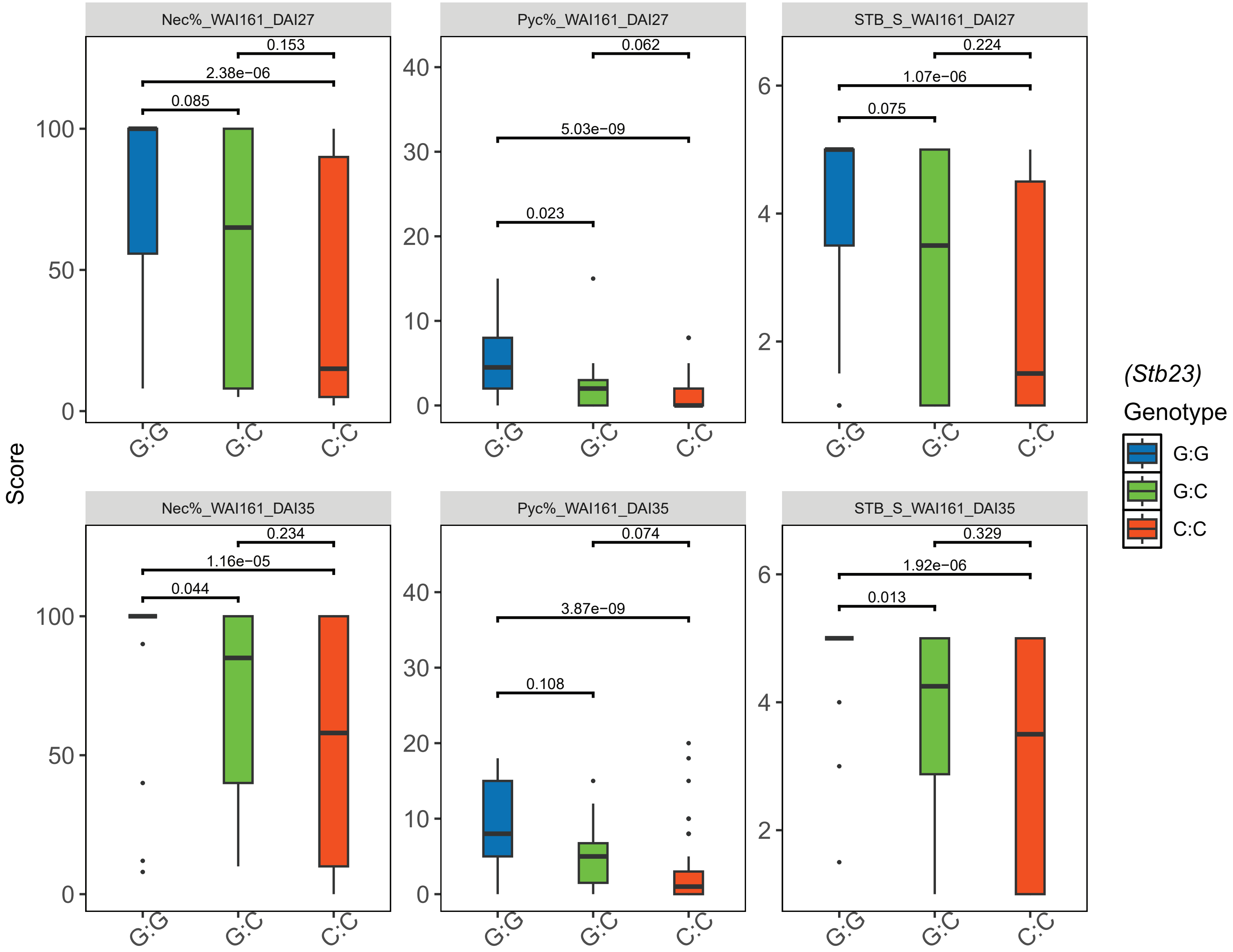


**Supplementary Fig. S13** Validation of the KASP marker snp_1D1217527 tightly linked to *Stb23* in segregating populations Comparisons of additive effects among homozygous and heterozygous lines in X6135 $F_4$ population using the isolate WAI161. Traits at the seedling stage include the percentage of necrotic leaf area (Nec%) on the infected leaves, the pycnidia density (Pyc%) in the necrotic leaf area, and the STB_S Scale 1 to 5. Traits were measured at 27 and 35 days after inoculation using the isolate WAI161. Red color represents homozygous resistant lines (C:C), green color represents the heterozygous lines (G:C), and blue color represents the homozygous susceptible lines (G:G). The p values acquired from multiple Wilcoxon test are shown above each pair of homozygous resistance and susceptible lines in each family.

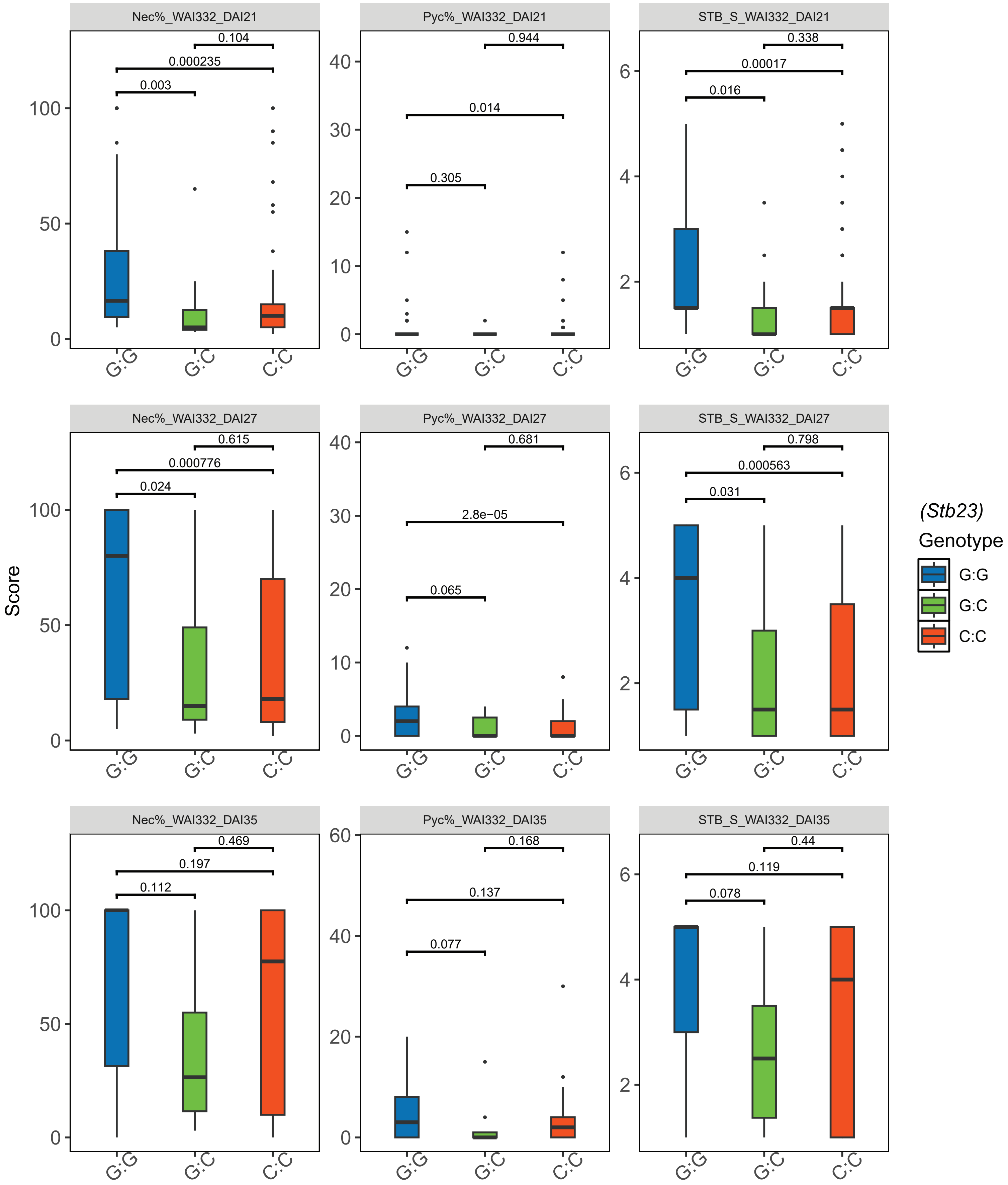


**Supplementary Fig. S14** Validation of the KASP marker snp_1D1217527 tightly linked to *Stb23* in segregating populations. Comparisons of additive effects among homozygous and heterozygous lines in X6135 $F_4$ population using the isolate WAI332. Traits at the seedling stage include the percentage of necrotic leaf area (Nec%) on the infected leaves, the pycnidia density (Pyc%) in the necrotic leaf area, and the STB_S Scale 1 to 5. Traits were measured at 21, 27 and 35 days after inoculation using the isolate WAI332. Red color represents homozygous resistant lines (C:C), green color represents the heterozygous lines (G:C), and blue color represents the homozygous susceptible lines (G:G). The p values acquired from multiple Wilcoxon test are shown above each pair of homozygous resistance and susceptible lines in each family.

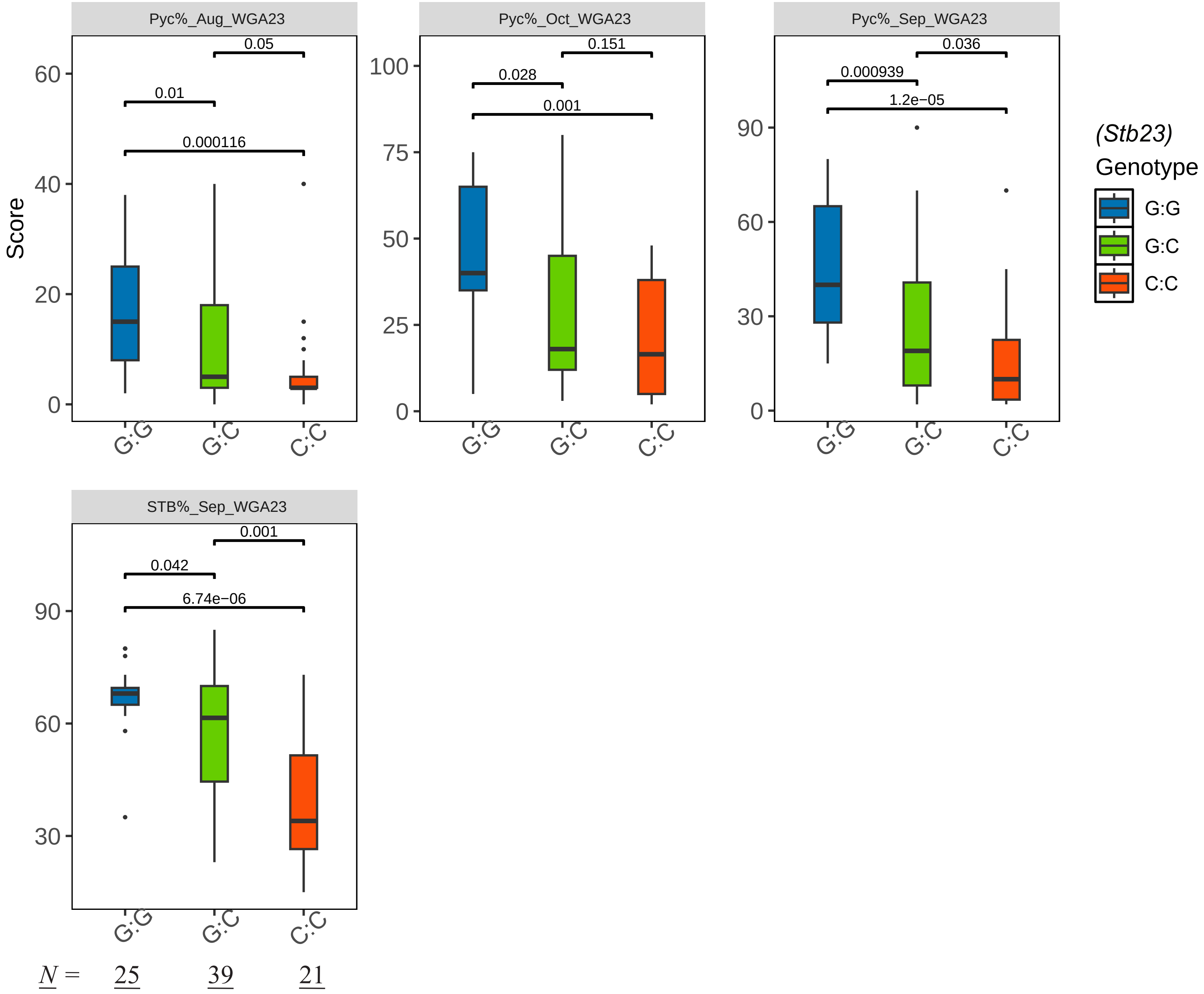


**Supplementary Fig. S15** Comparisons of additive effects of *Stb23* between homozygous and heterozygous lines in X6213 $F_2$ population for those traits collected from the field at Wagga Wagga NSW (WGA) in 2023. Traits at the adult-plant stage include the pycnidia density (Pyc%) in the necrotic leaf area and the percentage of STB infected leaf area on the whole plant (STB%) in August, September, and October. The number of individuals in each genotype category were shown at the bottom. Red color represents homozygous resistant lines (C:C), green color represents the heterozygous lines (G:C), and blue color represents the homozygous susceptible lines (G:G). The p values acquired from multiple Wilcoxon test are shown above each pair of homozygous resistance and susceptible lines in each family.

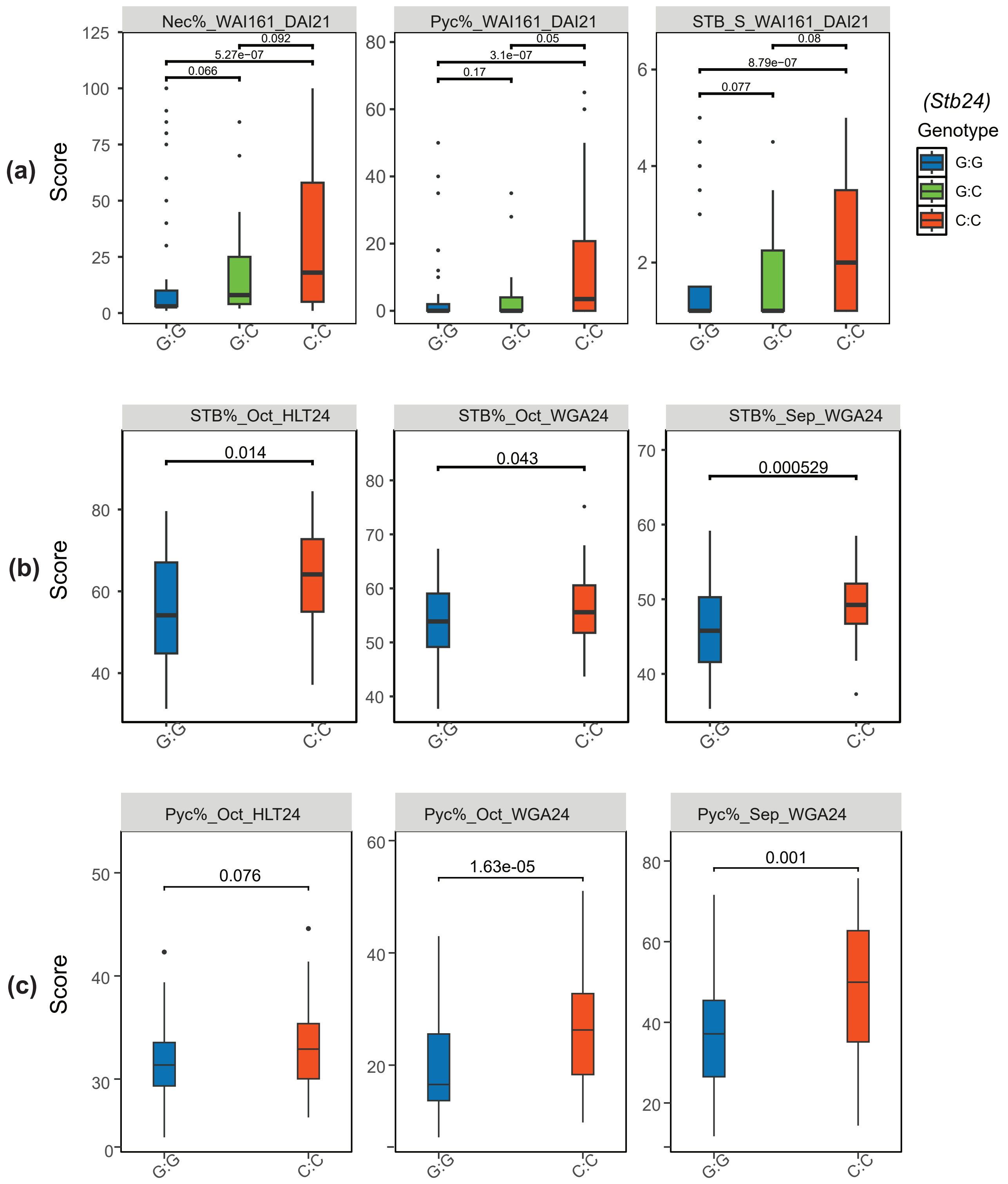


**Supplementary Fig. S16:** Validation of the KASP marker snp_3D1077880 tightly linked to *Stb24* in segregating population. **(a)** Comparisons of additive effects among homozygous and heterozygous lines in X6129 $F_4$ population using the isolate WAI161. Traits in the analysis include Nec%, Pyc%, and STB_S collected after 21 days after inoculations. **(b) and (c)** Comparisons of additive effects between homozygous resistant and susceptible lines in X6129 $F_6$ population for those traits collected from the field at Wagga Wagga NSW (WGA) and Hamilton Vic (HLT) in 2024. Traits in the analysis include the percentage of STB infected leaf area on the whole plant (STB%) and the pycnidia density (Pyc%) in the necrotic leaf area collected in September and October. Blue color represents homozygous resistant lines (G:G), green color represents the heterozygous lines (G:C), and red color represents the homozygous susceptible lines (C:C). The p values acquired from multiple Wilcoxon test are shown above each pair of homozygous resistance and susceptible lines in each family.

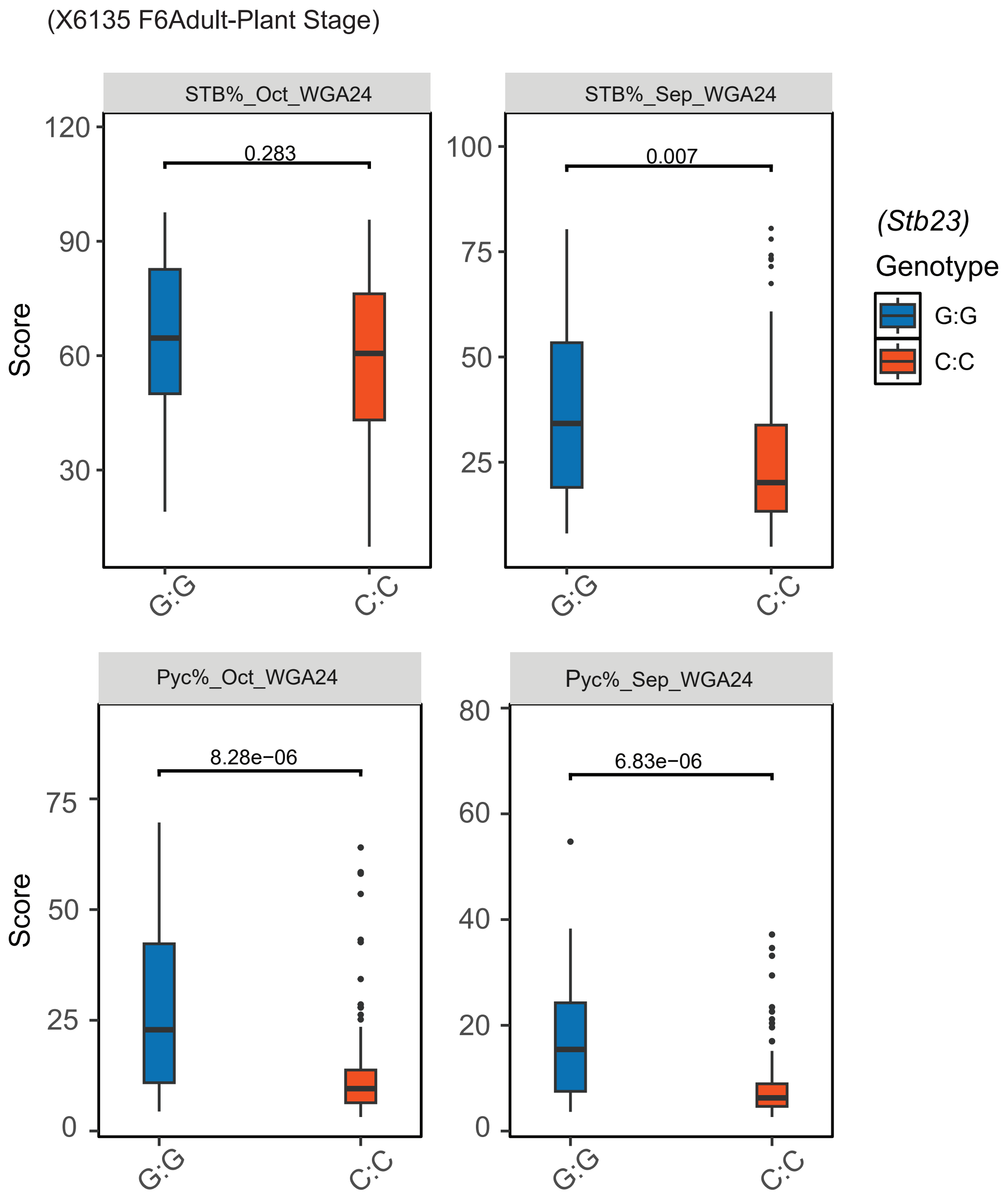


**Supplementary Fig. S17** Validation of the KASP marker snp_1D1217527 tightly linked to *Stb23* in segregating populations Comparisons of additive effects between homozygous resistant and susceptible lines in X6135 $F_6$ population for those traits collected from the field at Wagga Wagga NSW (WGA) in 2024. Traits in the analysis include the percentage of STB infected leaf area on the whole plant (STB%) and the pycnidia density (Pyc%) in the necrotic leaf area collected in September and October. Red color represents homozygous resistant lines (C:C) and blue color represents the homozygous susceptible lines (G:G). The p values acquired from multiple Wilcoxon test are shown above each pair of homozygous resistance and susceptible lines in each family.

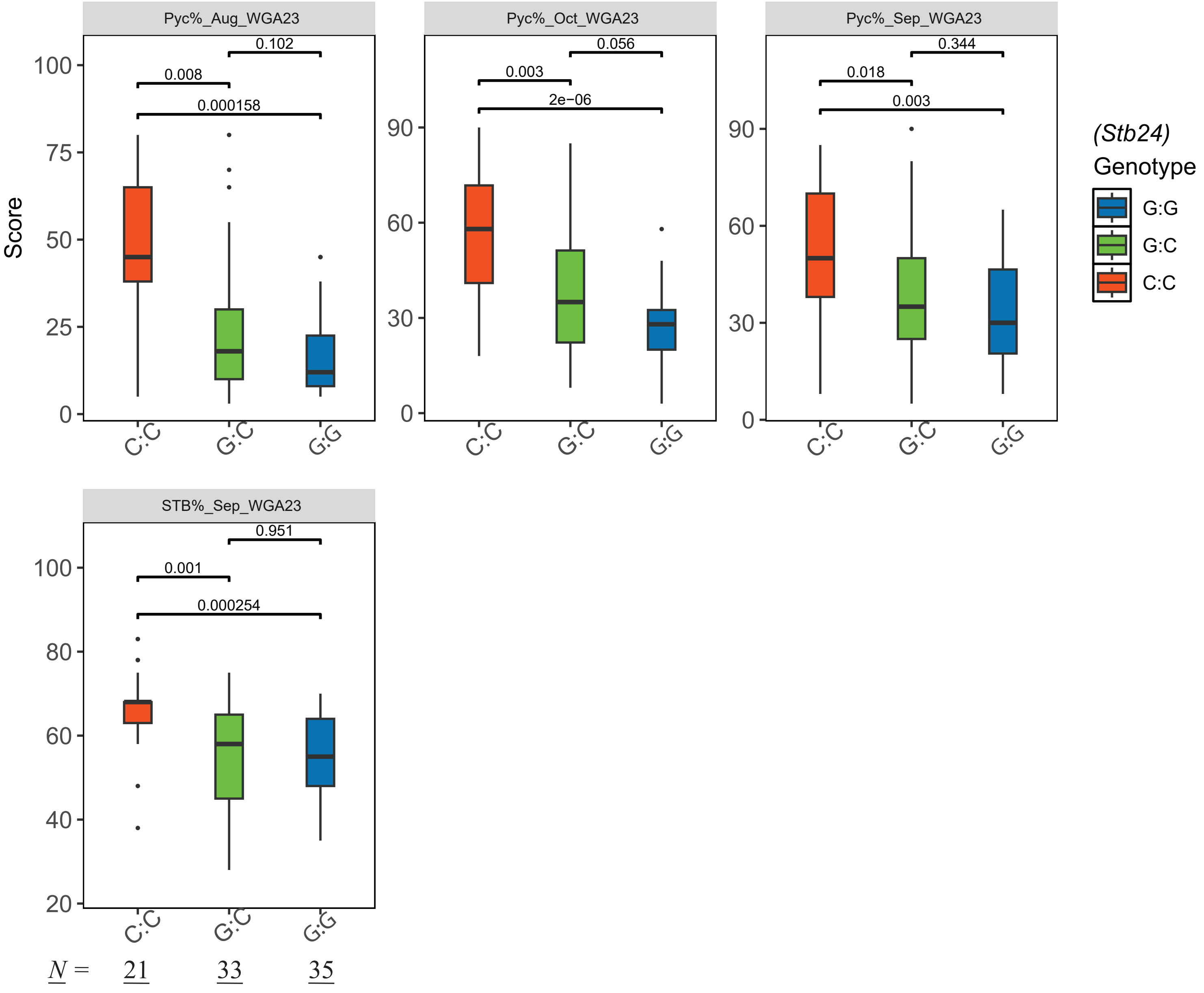


**Supplementary Fig. S18** Comparisons of additive effects of *Stb24* between homozygous and heterzygous lines in X6211 $F_2$ population for those traits collected from the field at Wagga Wagga NSW (WGA) in 2023. Traits at the adult-plant stage include the pycnidia density (Pyc%) in the necrotic leaf area and the percentage of STB infected leaf area on the whole plant (STB%) in August, September, and October. Blue color represents homozygous resistant lines (G:G), green color represents the heterozygous lines (G:C), and red color represents the homozygous susceptible lines (C:C). The p values acquired from multiple Wilcoxon test are shown above each pair of homozygous resistance and susceptible lines in each family.

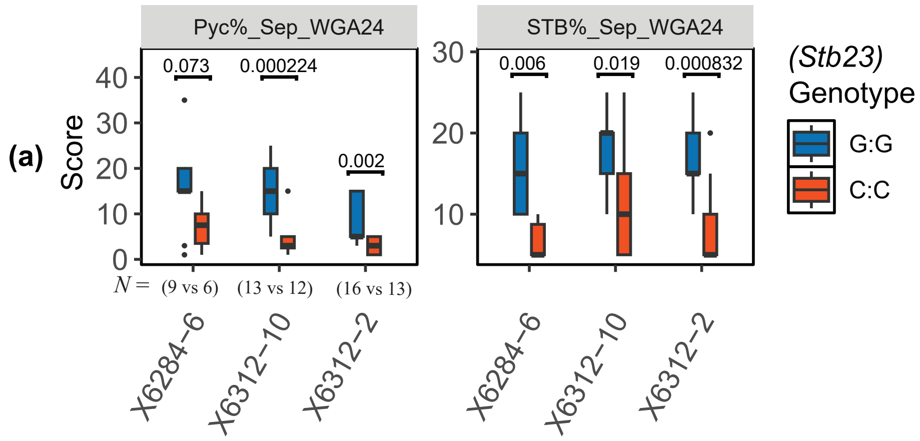


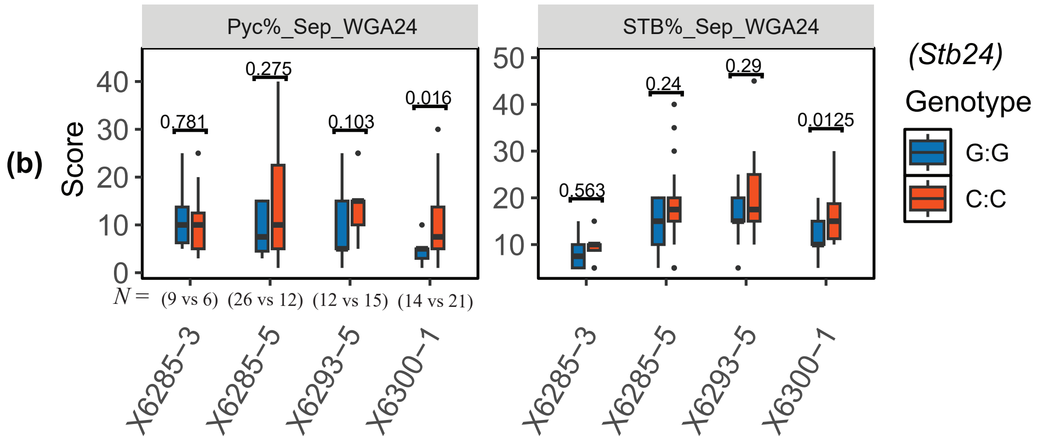


**Supplementary Fig. S19** Comparisons of additive effects between homozygous resistant lines and susceptible lines on *Stb23* and *Stb24.* **(a)** Comparisons of additive effects of *Stb23* among three different $F_2$ families X6284-6, X6312-10, and X6312-2. Red color represents homozygous resistant lines (C:C) and blue color represents the homozygous susceptible lines (G:G). **(b)** Comparisons of additive effects of *Stb24* among four different $F_2$ families, X6285-3, X6285-5, X6293-5, and X6300-1. Blue color represents homozygous resistant lines (G:G) and red color represents the homozygous susceptible lines (C:C). The number of homozygous lines used in the analysis shown under the bars. The p values acquired from multiple Wilcoxon test are shown above each pair of homozygous resistance and susceptible lines in each family.
